# Declarative Memory Through the Lens of Single-Trial Peaks in High-Frequency Power

**DOI:** 10.1101/2025.01.02.631123

**Authors:** Adam J.O. Dede, Zachariah R. Cross, Samantha M. Gray, Joseph P. Kelly, Qin Yin, Parisa Vahidi, Eishi Asano, Stephan U. Schuele, Joshua M. Rosenow, Joyce Y. Wu, Sandi K. Lam, Jeffrey S. Raskin, Jack J. Lin, Olivia Kim McManus, Shifteh Sattar, Ammar Shaikhouni, David King-Stephens, Peter B. Weber, Kenneth D. Laxer, Peter Brunner, Jarod L. Roland, Ignacio Saez, Fady Girgis, Robert T. Knight, Noa Ofen, Elizabeth L. Johnson

## Abstract

Declarative memory depends on the coordination of local processing, indexed by high-frequency broadband (HFB) activity, with global network organization, indexed by theta oscillations. However, theta and HFB exhibit asynchronous timing, raising the question of how results of local processing are communicated throughout the network. Using intracranial EEG in patients performing a recognition memory task, we examined this coordination across the medial temporal lobe (MTL) and prefrontal cortex (PFC). HFB peak activity was earlier in the MTL than PFC. Anchoring analyses of theta phase clustering and connectivity to HFB peaks revealed strong phase clustering locked to HFB peaks in the PFC, as well as connectivity between the PFC and MTL that predicted individual memory performance. Graph analysis revealed specific connections amidst sparse network connectivity during memory success. This study demonstrates that transient brain states linked to internal physiological events support memory and refines our understanding of local and network-level process interactions.

**Highlights:**

- Memory-linked theta activity is time-locked to internal brain events
- Network connectivity changes dynamically during memory processing
- Sparse network connectivity supports successful memory
- Specific sequences of transient states may be critical for declarative memory

## INTRODUCTION

High-frequency EEG components reflect local processing ^1–5^, while low-frequency components support network organization and information flow ^6^. The coordination of local and network-level signals is essential for complex processes like declarative memory, and both theta (2-8 Hz) and high-frequency broadband (70-150 Hz; HFB) signals have been extensively studied during memory tasks ^7–20^. Precisely how these signals interact remains unclear.

Utilizing intracranial EEG (iEEG), studies show that HFB activity peaks at different times across brain regions during memory encoding. Peak latencies progress along the ventral visual pathway to the medial temporal lobe (MTL) and later to association regions, including the prefrontal cortex (PFC) ^7–10,12^. PFC activity occurs earlier during retrieval ^7,8,11^. These findings suggest that complex processing occurs through sequences of transient states ^21–23^. By contrast, theta activity spans the brain shortly after stimulus onset ^9,24^, distinct from HFB activity ^10,13–20^. The question arises: if theta reflects network organization but does not coincide with local HFB processing, how are local results shared across the network?

The method of trial alignment may play a critical role in understanding this interaction. Peaks in HFB activity exhibit inter-trial variability ^25^, are brief ^26–28^, and resemble point processes similar to neuronal bursts ^29,30^. If theta activity were time-locked to HFB peaks, then trial-averaging to external events and long temporal epochs could obscure the true dynamics of these signals ^23^.

We hypothesized that theta network modulation would align with HFB peaks during successful memory encoding and retrieval. To test this, we analyzed iEEG data from epilepsy patients performing a memory task, focusing on five regions of interest (ROIs): hippocampus (Hip), parahippocampal gyrus (PHG), anterior cingulate cortex (ACC), dorsolateral prefrontal cortex (dlPFC), and polar prefrontal cortex (pPFC). The latency of peaks in HFB activity replicated the previously reported sequence of activity ^7,9,13^. We then examined how theta signal organization related to memory success, assessing both local phase clustering and between-region connectivity with time aligned either to local HFB peaks (HFB-locked) or to stimulus onset (image-locked).

HFB-locked analyses revealed mnemonic effects in theta phase clustering and connectivity, which were altered or absent in image-locked analyses. For instance, during encoding, connectivity between PFC and MTL regions was absent after stimulus onset in the image-locked analysis but present in the HFB-locked analysis, where it was significantly stronger for successful encoding and predicted individual memory performance. Additionally, graph-theoretic analysis revealed that connections between ROIs that increased with memory success were accompanied by a broad reduction in other network connections—a result that challenges prior models based on long-duration connectivity analyses ^31^. These results suggest that memory processing relies on a sequence of transient network states, characterized by coordinated changes in both high- and low-frequency signals.

## RESULTS

Thirty-six participants performed an image recognition memory task (Figure 1A) ^32–34^. The age, sex, and memory performance of participants did not vary by electrode coverage across five ROIs (Table 1; Figure 1B). To relate signals at lower frequencies to peak HFB activity, channels were selected within ROIs that exhibited task-reactive increases in HFB activity (see methods; Figure 1C) ^32,35–39^. The proportion of channels exhibiting reactivity varied between regions ( χ^2^(4) = 19, *p* =. 0009) (Figure 1D). All subsequent analyses utilized only channels identified as reactive.

**Figure 1.**
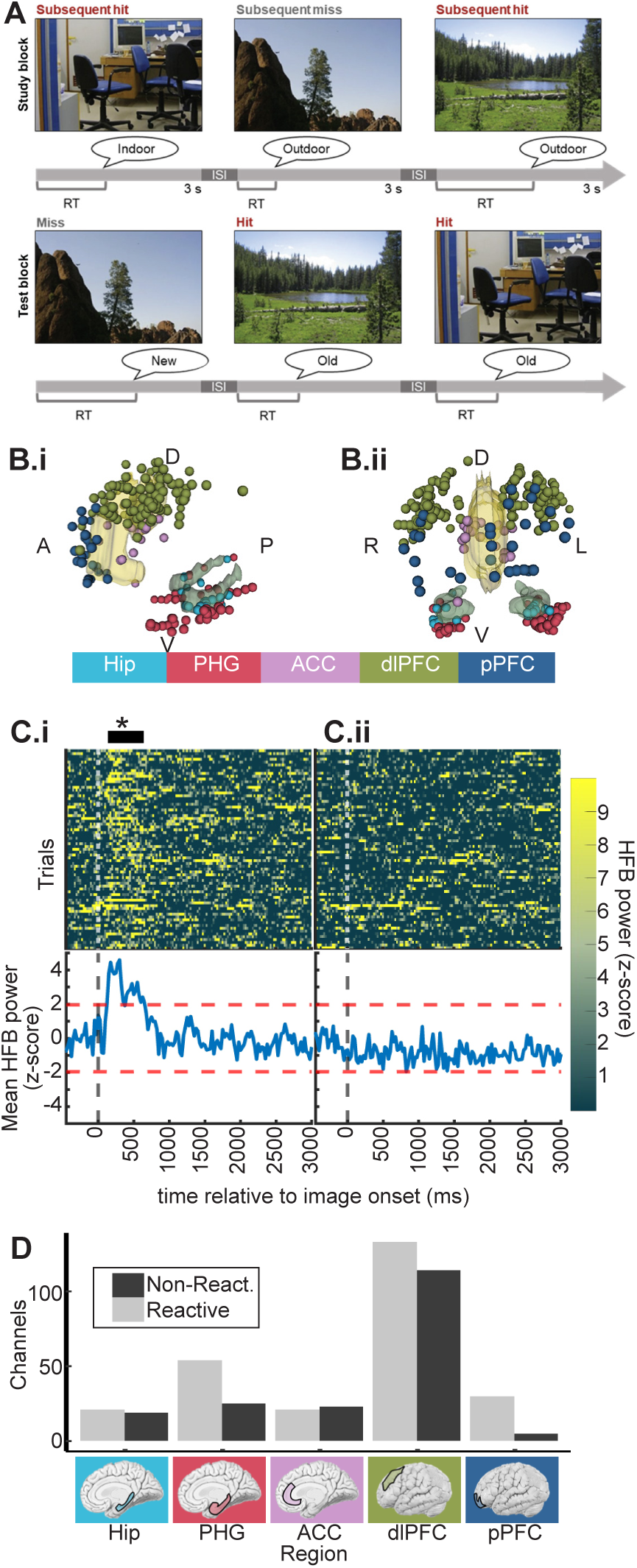
Task and channel coverage. **A.** Schematic of the behavioral task. During study blocks, participants made indoor/outdoor judgements while viewing images of natural scenes. During test blocks, participants made old/new judgements while viewing the same images of natural scenes intermixed with novel images. **B.** Channel coverage combined across participants. i. Lateral view. ii. Face-on view. ACC and Hip ROIs are shown with 3D models for reference. **C.** Example channels demonstrating detection of reactive channels. Top panels display heatmaps with trials on the y axis, time relative to image onset on the x axis, and the HFB power indicated by color. Bottom panels display the mean HFB power across trials on the y axis and time on the x axis. Notice that in panel i there is a response soon after image onset that exceeds the threshold of z>2 for more than 50 continuous ms (black horizontal line marked by asterisk above panel). By contrast, in panel ii, the mean response does not exceed the z-score threshold. Example channels were drawn from within a single participant’s PHG demonstrating within region heterogeneity. **D.** Summary of reactive and non-reactive channel counts across all ROIs. Notice that almost all pPFC channels were reactive. By contrast, other regions were more split.

**Table 1.** Participant demographics and key statistics. Age, memory accuracy, d’, and trial count columns are formatted as mean (SD).

| region | total channels | reactive channels | n (females) | age | memory accuracy | memory d' | hit trial count | miss trial count | ENCODING<br>low freq peak | ENCODING<br>med. freq peak | ENCODING<br>max ITPC | RETRIEVAL<br>low freq peak | RETRIEVAL<br>med. freq peak | RETRIEVAL<br>max ITPC |
| --- | --- | --- | --- | --- | --- | --- | --- | --- | --- | --- | --- | --- | --- | --- |
| Hip | 40 | 21 | 9 (3) | 19.01 (4.7) | .50 (.22) | 1.54 (.80) | 59.8 (36.0) | 24.3 (13.3) | 4.9 | - | 6.1 | 4.2 | 27.1 | - |
| PHG | 79 | 54 | 23 (9) | 18.4 (4.7) | .51 (.21) | 1.54 (.73) | 50.3 (24.5) | 21.2 (12.4) | 5.7 | - | 20.9 | 5.9 | - | 9.5 |
| ACC | 44 | 21 | 11 (3) | 19.4 (4.8) | .48 (.21) | 1.54 (.71) | 52.1 (37.9) | 22.8 (10.8) | 4.1 | - | 5.9 | 5.3 | 24.3 | - |
| dIPFC | 247 | 133 | 27 (9) | 18.7 (4.8) | .50 (.22) | 1.53 (.76) | 51.6 (28.2) | 21.0 (12.6) | 6.6 | - | 19.4 | 7.6 | 31.5 | 19.4 |
| pPFC | 35 | 30 | 8 (3) | 20.2 (4.32) | .40 (.19) | 1.16 (.63) | 41.5 (10.5) | 21.6 (6.4) | 2 | - | 6.1 | 4.9 | - | - |

### Latency of HFB activity replicates known sequence

Our main analysis focused on the latency of HFB peaks. A parallel analysis of HFB power is presented in Supplemental Figure 1.

We calculated the latency of HFB peak activity between image onset and behavioral reaction time (RT) for each trial on each channel (Figure 2A-C). Because peak latencies were detected between image onset and RT, they were biased to exhibit a relationship with RT (correlation between HFB peak latency and RT: *r* =. 45, *p* < 2*e* − 16, *t*(40281) = 102). That is, if it is the case that local HFB peaks form a sequence of activity, then this sequence may sometimes occur at a faster or slower rate without altering the sequence. This possibility is consistent with the observed HFB peak latency and RT correlation. Critically, if this is the case, then the sequence is not fundamentally altered on faster versus slower trials. To adjust for this, we divided peak latencies by RTs and observed no relationship between normalized HFB peak latency and RT (*r* =−. 008, *p* =. 1, *t*(40281) =− 1. 6). Thus, the relative timing of HFB peaks did not vary significantly with behavior.

**Figure 2.**
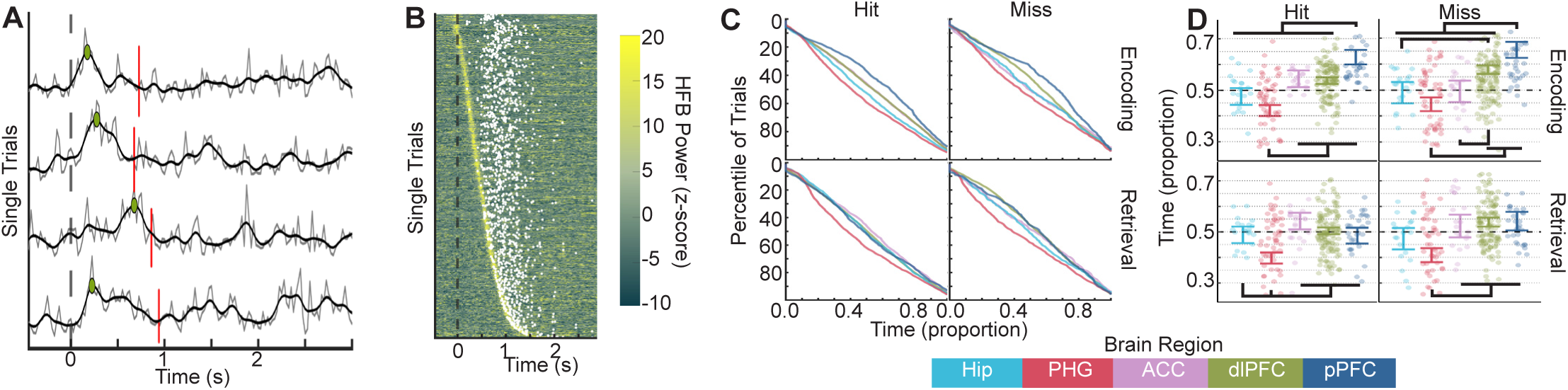
High-frequency broadband (HFB) peak latency timing changes with memory. **A**. In grey, four sequentially recorded examples of single trial HFB traces are displayed from a single channel. In black, the same traces are displayed after application of a Gaussian smoothing kernel. The broken vertical red line indicates the behavioral RT of each trial. The dashed vertical black line indicates image onset. The green dots indicate the peak HFB activity on each trial. Note the lack of consistency in peak latency. **B**. The heatmap displays single trial HFB time series from channels in the hippocampus during subsequent hit trials. Trials have been sorted by latency of the peak HFB power. White dots indicate the behavioral RT of each trial, which was a poor predictor of the latency of the HFB power peak. **C**. Each cumulative distribution plot displays the timing of the latency of peak HFB activity across all trials for subsequent hit (top left), subsequent misses (top right), retrieval hits (bottom left), and retrieval misses (bottom right). Each region’s trial distribution is shown with a different line. Because different numbers of trials were observed in different behavioral conditions and for different regions, trial is plotted as a percentile of trials on the y axis. Because reaction times were variable between individuals and conditions, time is plotted on the x axis as a proportion of time such that 0 is image onset and 1.0 is behavioral response. **D.** Each grouped scatter plot displays the mean time of peak HFB latency for each channel grouped by region. Time relative to image onset is displayed as a proportion on the y axis. Error bars display the 83% confidence interval around model estimates ^78,79^. For example, note that while the hippocampus (light blue) and parahippocampal gyrus (red) led the dlPFC and pPFC during successful encoding, these PFC regions were active simultaneously with the Hip during successful retrieval.

Linear mixed effects modeling of latency as a function of hit/miss by encode/retrieve by region revealed different temporal dynamics in different regions and as a function of mnemonic success during encoding versus retrieval (Figure 2D). Specifically, there were main effects of hit/miss (χ^2^ (1) = 17, *p* =. 00003), encode/retrieve (χ^2^ (1) = 48, *p* = 4*e* − 12) and region (χ^2^ (4) = 72, *p* = 8*e* − 15), and the interactions were significant between hit/miss and region (χ^2^ (4) = 14, *p* =. 007) and between encode/retrieve and region (χ^2^ (4) = 47, *p* = 2*e* − 9).

During successful encoding (hit trials), the PHG exhibited an HFB peak that occurred earlier than in all three PFC ROIs, with the pPFC latency occurring later than in all other ROIs. During failed encoding (miss trials), MTL ROIs were co-active with the ACC, which shifted earlier, and the dlPFC latency occurred later. HFB peaks in the ACC and PHG also exhibited mnemonic changes in power during encoding (Supplemental Figure 1C).

During retrieval of studied scenes (hit trials), the PHG exhibited the earliest peak, and the other four ROIs exhibited latencies similar to each other. This pattern was similar for misses, although both the dlPFC and pPFC exhibited significantly delayed latencies during miss trials relative to hits. Because it is possible that these timing effects were driven by individual differences between participants with coverage of different ROIs, we confirmed this analysis in several alternate ways (Supplemental Figure 2). These analyses yielded qualitatively similar results.

Taken together, these results replicate previous reports of the sequence of activity underlying declarative memory ^7,9,13^ and how it changes as a function of encoding/retrieval and memory success. Unlike the separation between MTL and PFC ROIs during successful encoding, HFB activity during hit retrieval trials still peaks first in the PHG, but then the Hip and PFC ROIs all exhibited similar peak latencies. When memory failed, these timing effects were altered.

### Mnemonic differences in theta phase clustering depend on time alignment

For the output resulting from local processing to be communicated to the broader network, we reasoned that theta oscillations would be systematically clustered around HFB peaks, reflecting organization of theta networks at the time of local HFB peaks. We compared encoding and retrieval trials sorted by memory outcome (subsequent hit, subsequent miss, hit, miss) using theta phase clustering (inter-trial phase clustering [ITPC]; see methods), a measure of the consistency of oscillatory phase at a particular time point across trials. This analysis was performed twice for each ROI: HFB-locked and image-locked. In this way, it was possible to compare how signals were organized relative to the internal event of peak local activity versus the external event of stimulus presentation. Because we were focused on fine-grained temporal organization of low-frequency oscillations, phase-based measurements were of primary interest. A parallel analysis of low-frequency power is presented in Supplemental Figure 3.

In the HFB-locked analysis, phase clustering was stronger for subsequent hit and hit trials over subsequent miss and miss trials (Figure 3A). Specifically, we observed greater phase clustering for subsequent-hit compared to subsequent-miss trials across frequencies concentrated in the theta range in all five ROIs, as well as greater phase clustering for hit compared to miss trials at retrieval in the PHG and dlPFC (all cluster-corrected p<.05). These results indicate that increased phase clustering time-locked to HFB peaks is an important organizing principle of signals associated with successful memory. For image-locked ITPC analyses, there appeared to be a gradient across ROIs such that the MTL ROIs exhibited strong phase clustering across 2 to 10 Hz between -75 and 825 ms that differed between (subsequent) hit and miss trials, while this effect became weaker in the ACC, yet weaker in the dlPFC, and non-existent in the pPFC (Figure 3B).

**Figure 3.**
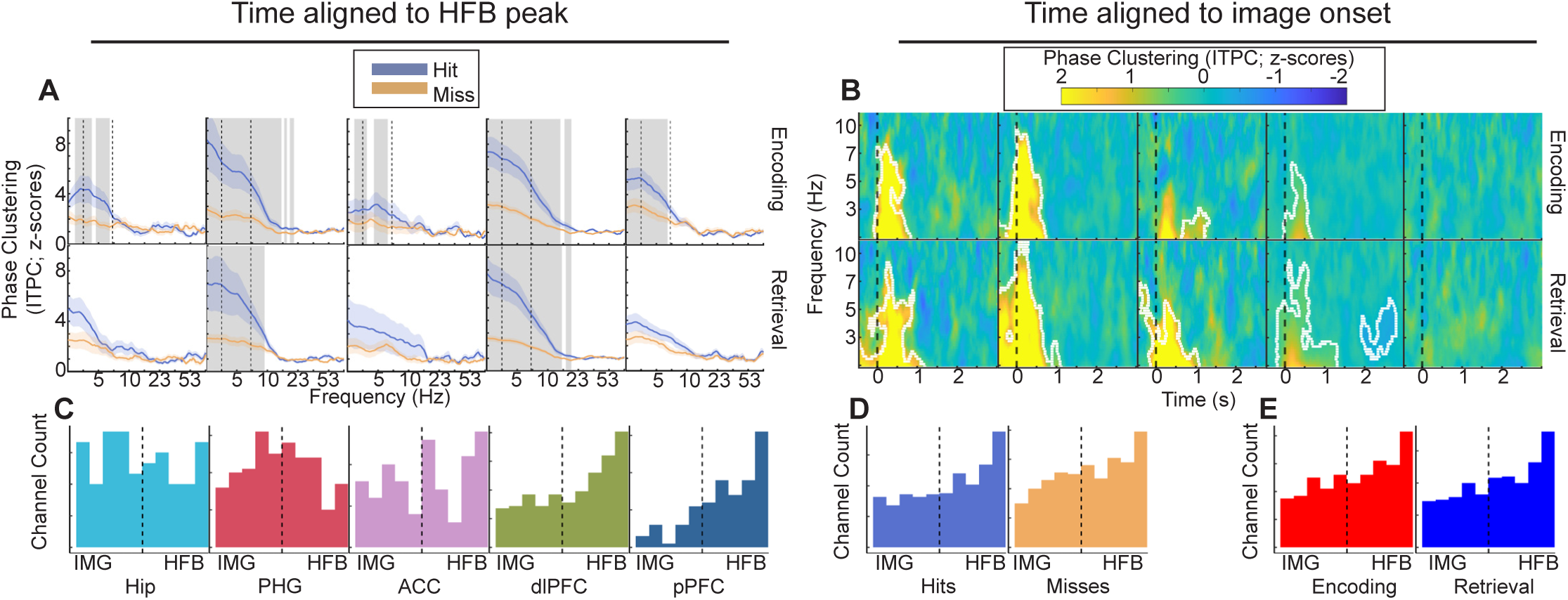
Patterns of mnemonic differences in phase clustering are dependent on time alignment. **A**. Line plots display inter-trial phase clustering (ITPC) spectra aligned to the time point of the HFB peak. For all panels, the x-axis displays frequency, and the y-axis displays ITPC in z-scored units. Encoding and retrieval data are plotted along the top and bottom rows, respectively. Orange and blue lines are the average spectra of (subsequent) hit and miss trials, respectively. Vertical gray shaded regions indicate p<.05 for the difference between hit and miss spectra after cluster correction. Colored shaded regions indicate the standard error of the mean. **B**. Heatmaps display the mean difference in z-scored ITPC between hit and miss trials. The x-axis displays time in seconds relative to image onset, and the y-axis displays frequency. Color indicates the difference between (subsequent) hit and miss trials in units of z-scored ITPC. Encoding and retrieval data are plotted along the top and bottom rows, respectively. Note that the dlPFC and pPFC exhibited strong hit/miss phase clustering effects when analysis was locked to local HFB peaks (panel A), but phase clustering effects are weaker when analysis is locked to image onset. **C**. Histograms display the IMG-HFB index for different regions collapsed across hit/miss and encode/retrieve trial types. Values at the extremes indicated that phase clustering was observed exclusively when data were analyzed relative to image onset (left) or HFB peak (right). Intermediate values (near the vertical dashed line) indicated that the two treatments of time yielded similar results. Note that the distribution of channels in the dlPFC and pPFC is strongly skewed to the HFB end of the axis. **D**. The same as panel C but with histograms representing hit and miss trials collapsed across region and encoding/retrieval. **E**. The same as panel C but with histograms representing encoding and retrieval trials collapsed across region and hit/miss. The high channel count in the dlPFC leads to this region dominating the distribution shape in panels D and E, but the key observation is that IMG-HFB index values do not vary with hit/miss or encode/retrieve.

To quantify the relative strength of image-versus HFB-locked ITPC, we calculated a normalized IMG-HFB index score for each channel (see methods; Figure 3C-E). This score could range from -1 to +1 where +1 indicated that phase clustering was much stronger for the HFB-locked over the image-locked analysis. When IMG-HFB scores were modeled as a function of hit/miss, encode/retrieve, and region, only the main effect of region was significant (χ^2^ (4) = 31, *p* = 3*e* − 6). Visual inspection of Figure 3C revealed that the dlPFC and pPFC exhibited stronger HFB-locked phase clustering, whereas IMG-HFB scores in other regions were more balanced.

Considering both mnemonic differences and the image-versus HFB-locked relative strength of phase clustering, these results suggest that the Hip participates more strongly in stimulus-driven mnemonic processing, exhibiting stronger hit/miss differences in phase clustering in the image-locked analysis. By contrast, processing in the dlPFC and pPFC appears to be organized around internal events. The PHG and ACC appear to participate in both. Interestingly, although the Hip and PHG exhibited strong phase clustering in both image- and HFB-locked analyses, these regions exhibited markedly higher phase angle consistency across channels in the HFB-locked analysis (compare Supplemental Figure 4A to B). This implies that network organization across channels was more consistent when data were aligned with HFB peaks than when data were aligned with image onset.

### Mnemonic differences in theta connectivity depend on time alignment

To understand how connectivity between ROIs at low frequencies varied as a function of memory, we compared connectivity (pairwise phase consistency [PPC]; see methods)^40^ at encoding and retrieval trials sorted by memory outcome independently for each pair of channels. As with phase clustering, this analysis was performed with both image- and HFB-locked data. This analysis was sensitive to the fine time scale organization of low-frequency networks because our measure of connectivity is based on the consistency of the phase offset between channels at a particular time point across trials.

Many pairwise connections differed (p<.05) after cluster correction based on memory outcome (hereafter termed mnemonic connections), but these were not evenly distributed across encoding/retrieval, frequency, time, region, or analysis approach. In general, there were more mnemonic connections during retrieval than during encoding, and 72% of all mnemonic connections were increases in connectivity for positive memory outcomes (subsequent hits, hits) over negative memory outcomes (subsequent misses, misses). We focus on these positive mnemonic connections; see Supplemental Figure 5 for a schematic representation of negative mnemonic connections. Mnemonic connections were concentrated in frequencies below 10 Hz (Figures 4A,C), with encoding effects largely below 3.5 Hz and retrieval effects ranging up to 10 Hz. The timing of mnemonic connections varied between encoding and retrieval. Specifically, HFB-locked mnemonic connections occurred prior to HFB peaks during encoding, but were centered around HFB peaks during retrieval (Figure 4B). Image-locked mnemonic connections were most prevalent around image onset and late in the trial during encoding, but were observed across the entire trial during retrieval (Figure 4D). For visualization, mnemonic connections are indicated schematically in Figure 4E,F; time-frequency representations of connectivity are presented in Supplemental Figures 6-9.

**Figure 4.**
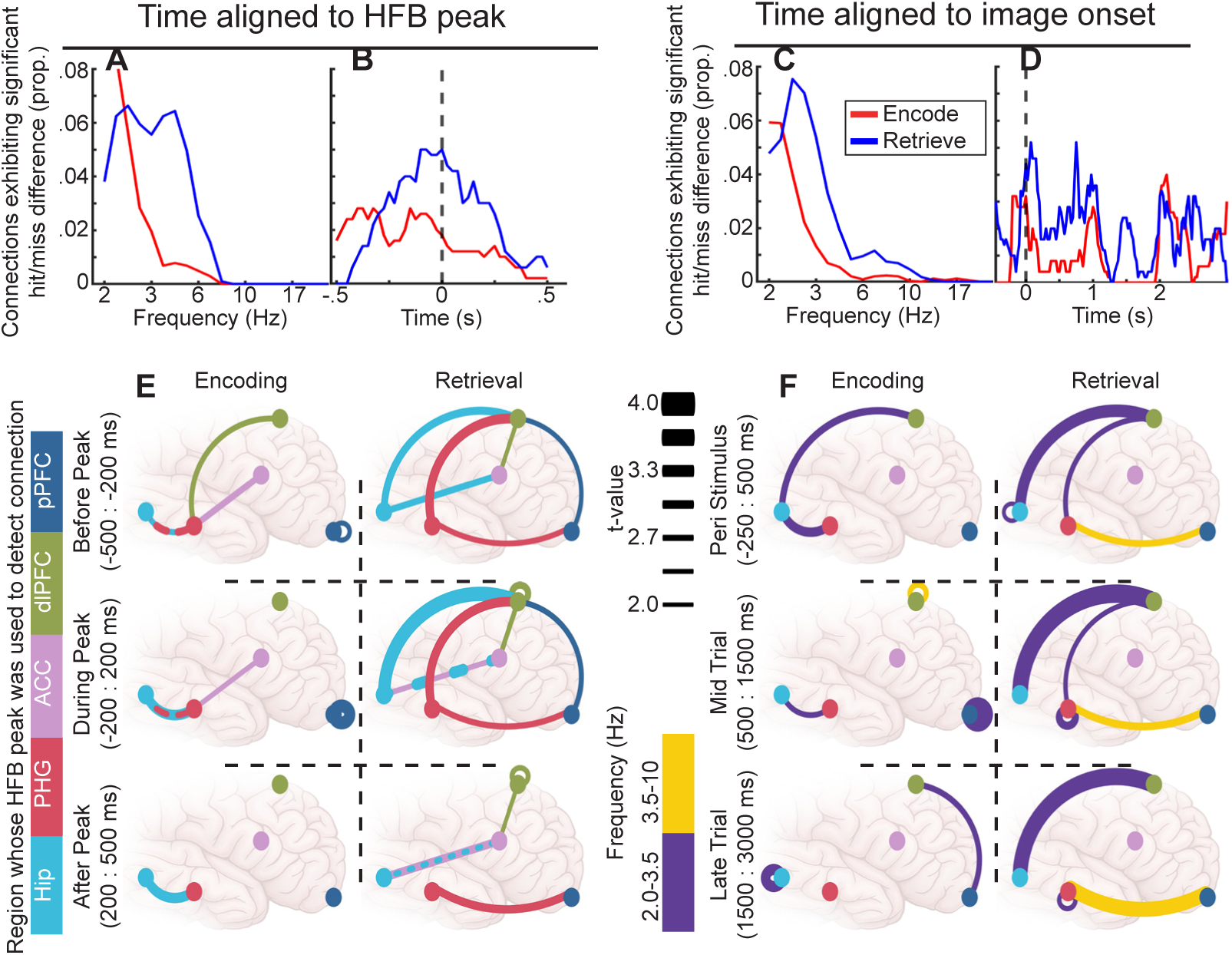
Patterns of connectivity were different depending on time alignment. In panels A-D the Y axes display the proportion of the time-frequency space for all tested connections that exhibited a significant difference between hit and miss trials. **A.** The x axis displays frequency. **B.** The x axis displays time relative to HFB peak. **C.** As A for image-locked analysis. **D.** The x axis displays time relative to image onset. **E.** The connection schematics display connections that were significantly increased for hit relative to miss trials for HFB-locked analysis. Line colors indicate the region whose HFB peak was used for temporal alignment. The rows display those connections that exhibited significance before (top), during (middle), and after (bottom) the HFB peak. The columns correspond to encoding (left) and retrieval (right). Dots represent color-coded region locations. The thickness of lines connecting dots represents the t-value of the difference between hit and miss trial connectivity. Connections between pairs of channels within a single ROI are represented with self-referencing loops. **F.** Line colors indicate the mean frequency of the significant difference between hit and miss trials. The connection schematics reflect mnemonic connections from the image-locked analysis. All other conventions are similar to E except that time is divided into peri stimulus (top), mid-trial (middle), and late-trial (bottom) periods. For hit less than miss differences, see Supplemental Figure 7.

HFB-locked analysis of encoding revealed mnemonic connections between the Hip and PHG, dlPFC and PHG, and ACC and PHG (Figure 4E). These pairwise connections were detected using data aligned in time to the HFB peak in one ROI. Consideration of which ROI’s HFB peak had been used to detect each mnemonic connection, combined with the timing of HFB peaks (see Figure 2D), suggested a dynamic sequence of connectivity patterns. Specifically, successful encoding was marked by connectivity within the MTL that spanned local HFB peaks, followed by connectivity between the PHG and dlPFC/ACC that preceded the HFB peaks in PFC ROIs. Analysis of retrieval revealed mnemonic connections between the MTL ROIs and all three PFC ROIs, but not within the MTL. Mnemonic connections were associated with the timing of PFC HFB peaks during encoding and with the timing of MTL peaks during retrieval, suggesting a reversal of connectivity direction.

Image-locked analysis of encoding revealed mnemonic connections between the Hip and dlPFC prior to image onset, within the MTL throughout the trial, and between the pPFC and dlPFC late in the trial (Figure 4F). Notably absent were connections between the MTL and PFC after image onset. Analysis of retrieval revealed a more extensive set of mnemonic connections. In particular, connections between the PFC and MTL were observed throughout the trial. Mnemonic connections involving the ACC were absent in image-locked analyses.

Taken together, mnemonic connections between MTL and PFC ROIs were more prevalent during retrieval than encoding. The MTL-PFC connections that were identified during encoding were aligned to the HFB peaks of PFC ROIs, suggesting that these late connections may be recruited by local activity in the PFC. By contrast, mnemonic connections during retrieval were more likely to be aligned to the HFB peaks of MTL ROIs, or in the case of image-locked connections, persisted throughout the trial. These results, together with temporal alignment between Hip and PFC HFB peak latencies (see Figure 2D), suggest early integration between the MTL and PFC to facilitate retrieval. Finally, mnemonic connections involving the ACC were only detectable in the HFB-locked analysis for both encoding and retrieval.

### Mnemonic increases in theta connectivity are contextualized by sparse network states

Most mnemonic changes were hit>miss as opposed to miss>hit differences (compare Figure 4E,F to Supplemental Figure 5). Were these connectivity increases specific or were connections throughout the brain strengthened at the same time as the connections detected between ROIs? To answer this question, for each significant change in ROI-ROI connectivity, each participant’s full channel X channel connectivity matrix was extracted for the time and frequency ranges of the significant difference. This was done separately for different trial types (subsequent hit, subsequent miss, hit, miss). For example, in the HFB-locked analysis of retrieval, hit trials were characterized by a significant increase in connectivity between the Hip and ACC concentrated in low frequencies after the ACC HFB peak (Supplemental Figure 9). All six participants with electrodes in both of these regions exhibited this difference (Supplemental Figure 10A). The connectivity matrices averaged across significant time and frequency ranges for a single representative participant were binarized and plotted in MNI anatomical space for visualization (Figure 5A.i,A.ii). Although the time and frequency ranges used to extract these values were chosen because of greater hit-over-miss connectivity between ROIs, visual inspection suggested that the network state across all channels was one of greater connectivity during miss trials. By contrast, when connectivity values were extracted at the same frequencies but averaged across a larger time window (between 0 and 1.5 seconds after image onset ^41^), there was little difference between the connectivity patterns observed for hit compared to miss trials (Figure 5A.iii, A.iv). Thus, for this example participant, the transient increase in Hip-ACC connectivity observed during successful retrieval appears to have been specifically important for memory success as opposed to being a random sample from an overall more connected network. The contrast of increased connectivity between ROIs with decreased connectivity across the brain may be a marker of increased network tuning to facilitate memory success.

**Figure 5.**
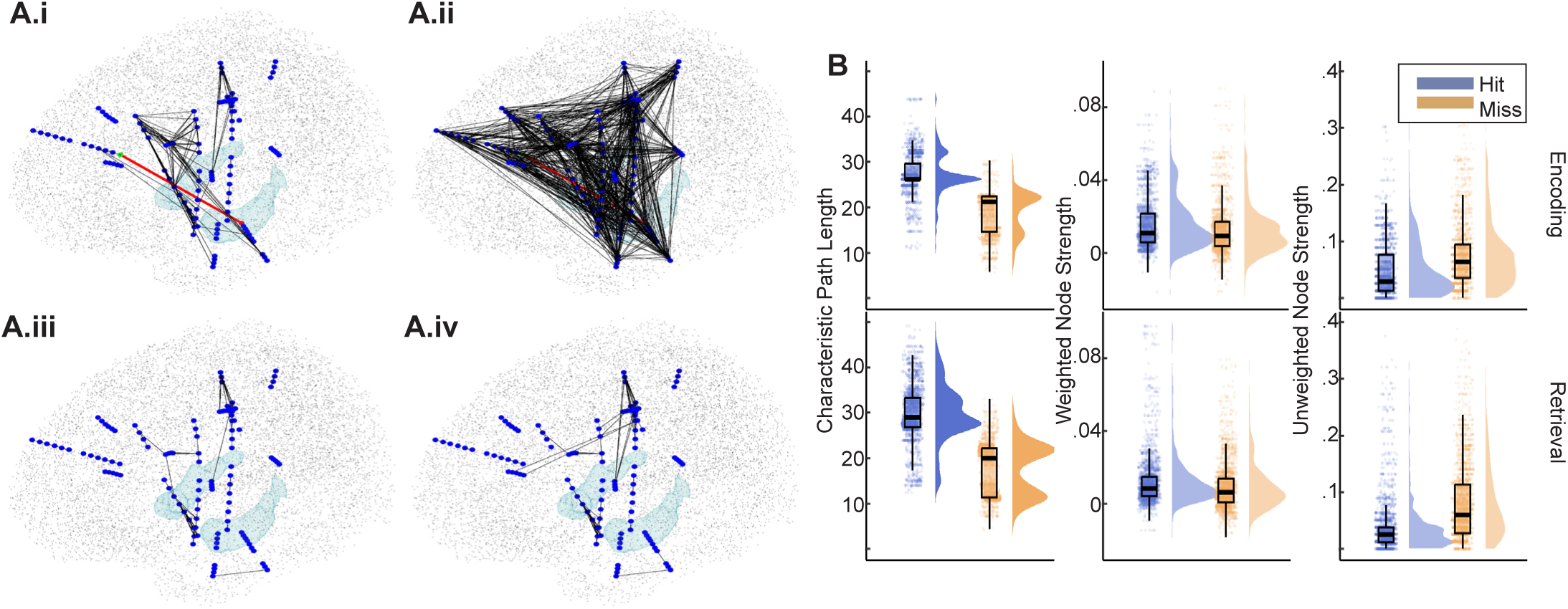
Short periods of increased connectivity between ROIs are characterized by a sparse overall network state. **A.** Plots display binarized connectivity graphs for a single participant between 2.6 and 6.6 Hz during retrieval. Blue dots indicate recording channels. Light blue surfaces indicate the location of the hippocampi for anatomical reference. Black stippling indicates the cortical surface. Black lines represent connections with connectivity strength (pairwise phase consistency) greater than .1. i and ii display connections averaged across the time window between -425 and -75 ms relative to the Hip HFB peak. The red line indicates a connection between the ACC and Hip, which was elevated during successful retrieval for all participants (Supplemental Figure 10A). i displays connections during hit trials. ii. displays connections during miss trials. Note that there are many more connections with PPC > .1 during miss trials. iii and iv display similar connection graphs with connectivity values averaged across the time window between 0 and 1500 ms relative to image onset. Note that far fewer connections with connectivity > .1 were found, and there is little difference between hit and miss trials. **B.** Graph theoretic measures taken for all connectivity graphs calculated using the time-frequency parameters of significant hit v. miss clusters. Characteristic path length and unweighted node strength both indicated more connected graphs during miss trials than hit trials, extending the effects seen in A across all participants and all connections.

To evaluate network tuning across all participants and all connections, graph theoretic analysis was applied to the channel X channel connectivity matrices for each participant at the time and frequency range of each significant change in connectivity detected between ROIs. Graph theory provides descriptive measures capable of quantifying the overall connectivity state of a network ^42^. Each significant change in connectivity was characterized by a particular analysis timing (HFB-locked vs. image-locked), time, frequency, region 1, and region 2. For statistical analysis, graph measures were combined across levels of time, frequency, and region 2. Graph measures were analyzed using linear mixed effects modeling as a function of memory outcome (subsequent hit/subsequent miss, or hit/miss) by region 1 by analysis timing. These analyses were repeated for encoding and retrieval separately. Thus, this analysis asked, what was the overall network state associated with mnemonic connections detected for each region and analysis timing separately? Three graph measures were calculated ^42^. The characteristic path length is the average distance needed to traverse the graph between any two channels. The weighted strength is the summed connectivity values across all channels paired with a single seed channel in region 1. The unweighted strength is the proportion of values greater than .1 across all channels paired with a single seed channel in region 1.

Across encoding and retrieval, there were main effects of memory outcome such that characteristic path length was longer and unweighted strength was lower during hit relative to miss trials (χ^2^ (1) > 788, *maximum p* < 2*e −* 16), (Figure 5B). Although weighted strength was higher for hit than for miss trials (χ^2^ (1) > 113, *p* < 2*e −* 16), the size of this effect was smaller than observed for the other two measures. These results echo the intuition gained from visual examination of Figures 5A.i and A.ii. That is, when specific ROI-ROI connections increased for successful memory outcome, the network more specifically tuned, resulting in longer characteristic path lengths, and the proportion of connections exhibiting high strength decreased (see Supplemental Figure 10 for results separated by ROI).

Taken together, these results demonstrate that increased connectivity between ROIs in association with successful memory was accompanied by decreased connectivity throughout the brain in general. Specific connectivity effects reflected transient and discrete temporal epochs.

### Phase clustering and connectivity within and between the MTL and dlPFC predict individual memory performance

Lastly, we conducted an exploratory analysis to correlate each of the 120 significant differences between trials classified based on memory outcome with individual memory performance (Figure 6). Specifically, for each participant, the mean mnemonic difference (subsequent hit vs. subsequent miss, or hit vs. miss) was calculated across channels per ROI for each signal component at encoding or retrieval. These participant-level values were then correlated with individual memory performance (d’). To ensure that results were not driven by single participants, correlations are only reported here if they were significant at the p<.05 level and maintained at p<.10 with the removal of any one participant. During both encoding and retrieval, phase clustering in the Hip and dlPFC (see Figure 3A-B) and connectivity between the dlPFC and PHG (see Figure 4E-F) predicted individual memory performance. Intriguingly, whereas phase clustering in the dlPFC and connectivity between dlPFC-PHG that predicted memory performance were locked to the HFB peak of the dlPFC during encoding, they were locked to image onset during retrieval. This further emphasizes the late, internally-driven role of the dlPFC during encoding versus its earlier role during retrieval.

**Figure 6.**
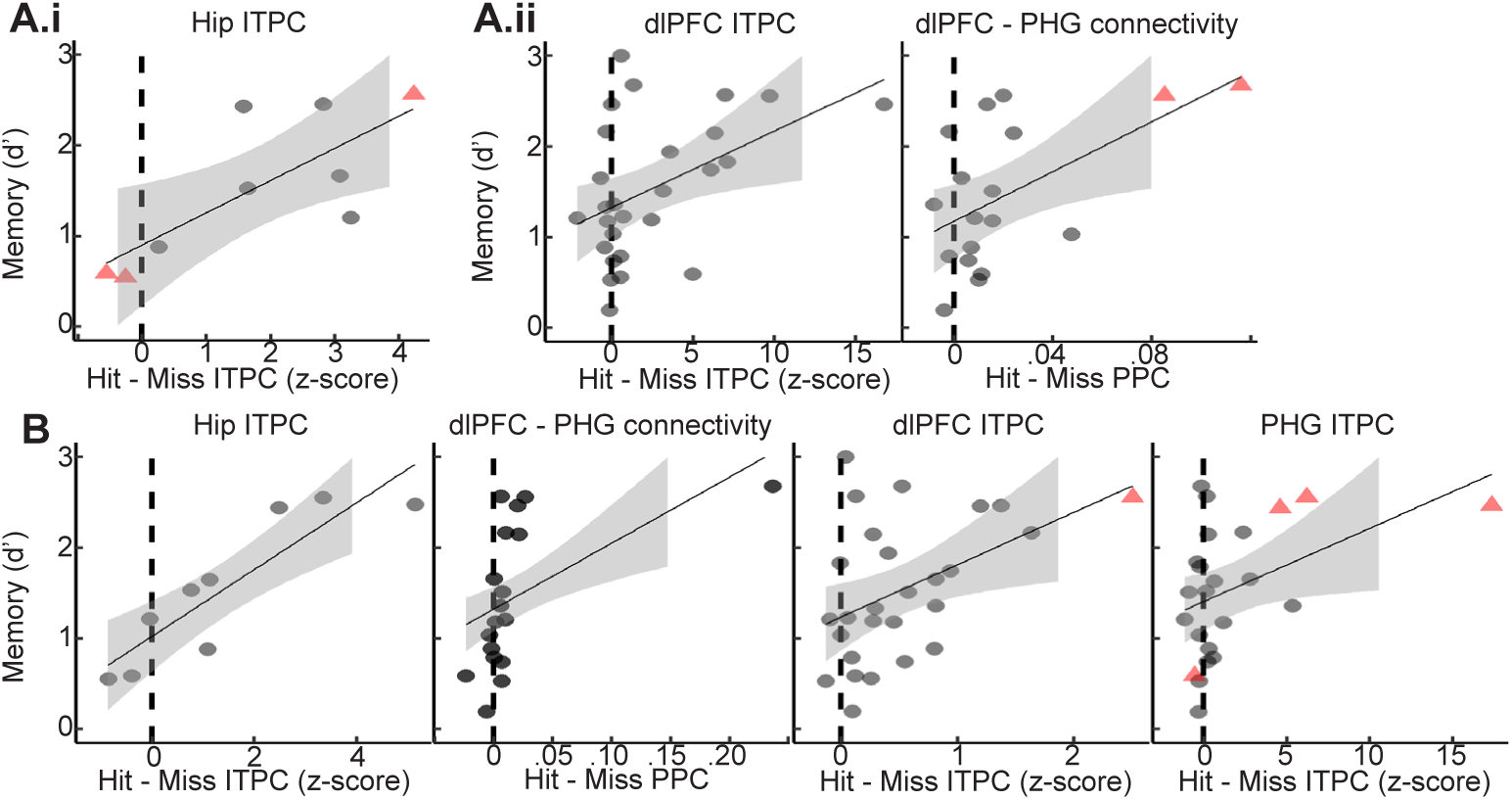
Individual memory performance was predicted by a subset of iEEG signal components. All scatter plots display significant relationships between individual-level signal components and memory performance. The x-axis displays the difference between the value of the signal component on hit minus miss trials. The y-axis represents behavioral memory performance (d’). Points represent means across channels within each participant. Red triangles indicate participants whose removal would result in the p value rising from below .05 to between .05 and .10. Relationships for which removal of any single participant caused the p value to rise above .1 are not shown. The signal component displayed along the x axis of each plot is indicated by its title. **A.i** Signal component aligned to image onset during encoding that predicted memory performance **A.ii** Signal components aligned to HFB peaks during encoding that predicted memory performance. **B.** Signal components aligned to image onset during retrieval that predicted memory performance. Note that ITPC at image onset, dlPFC ITPC, and dlPFC-PHG connectivity were important at both encoding and retrieval.

## DISCUSSION

The present results reveal different sequences of transient states associated with successful memory during encoding versus retrieval across a distributed network of MTL and PFC regions. To characterize these states, the latencies of HFB peaks served as key temporal anchors. Indeed, we replicated previous reports indicating that during memory encoding, HFB power peaks in MTL regions prior to PFC regions ^9,10,12,13,43^, and that there is earlier PFC involvement during successful compared to failed retrieval ^7^ (Figure 2D). To better understand the sequence of brain states that accompanied these HFB peaks, all other analyses were performed twice: with time aligned to HFB peak latencies and with time aligned to image onset. This analysis was designed to address a seeming paradox in prior literature. Specifically, if theta connectivity is a primary signal of network organization during cognitive task performance, then why does it seem not to coincide with local processing reflected in HFB activity?

A key idea of our dual approach was that many aspects of brain activity are internally organized rather than directly responsive to external stimulus events ^21,44^. Indeed, many effects were not detectable using a typical outside-in approach. This is not to say that analyzing data relative to internal brain events is superior. Five of the 7 robust correlations we identified between individual physiology and memory performance were found using metrics aligned to image onset (Figure 6). Rather, the present results emphasize that the use of internal timing markers yields complementary results compared to those obtained using external reference--particularly for events more distant from stimulus processing. For example, there were no mnemonic connections involving the ACC when data were aligned to image onset; ACC connections were revealed when data were aligned to HFB peaks, echoing rodent findings indicating this region as an important node in mnemonic processing ^45–47^. More generally, signals recorded throughout the PFC exhibited stronger phase clustering in HFB-locked than in image-locked analyses (Figure 3E). By contrast, signals in the MTL exhibited similar phase resetting across analytic approaches.

Across the signal components considered here, we found 120 significant differences between (subsequent) hit and miss trials (plus 27 related to power changes reported in Supplemental Materials). These differences were associated with transient time windows. Using these time windows combined with whether each effect was HFB-locked or image-locked, it is possible to integrate all of our results into two broad states for encoding and retrieval. When the results are integrated in this way, links to well-known cognitive phenomena become apparent.

Mean HFB peak latencies occurred early after stimulus onset in both MTL regions (Figure 7; left side, top panel). At this time, successful encoding was characterized by theta phase clustering in both regions and theta connectivity between these regions that was locked to their HFB peaks. Taken together, these physiological effects may represent the transformation of perceptual information into an enduring engram, a process for which the integrity of the MTL is critical and which may be indexed by HFB activity in these regions ^29,48^. Intriguingly, the HFB-locked increase in connectivity between MTL regions occurred prior to the PHG’s (earlier) HFB peak, but straddled the Hip’s (later) HFB peak (Supplemental Figure 7C), implying two separate phases of connectivity and echoing earlier findings of complex temporal dynamics within regions of the MTL ^49^. These mnemonic changes in the temporal organization of theta phase included the prediction of individual memory performance by Hip phase clustering (Figure 6). Thus, the picture emerges of a Hip-led MTL theta network. Indeed, we and others ^16,17,50–52^ have also observed Hip theta power increases prior to stimulus onset during successful encoding (Supplemental Figure 3F), emphasizing the early involvement of this structure.

**Figure 7.**
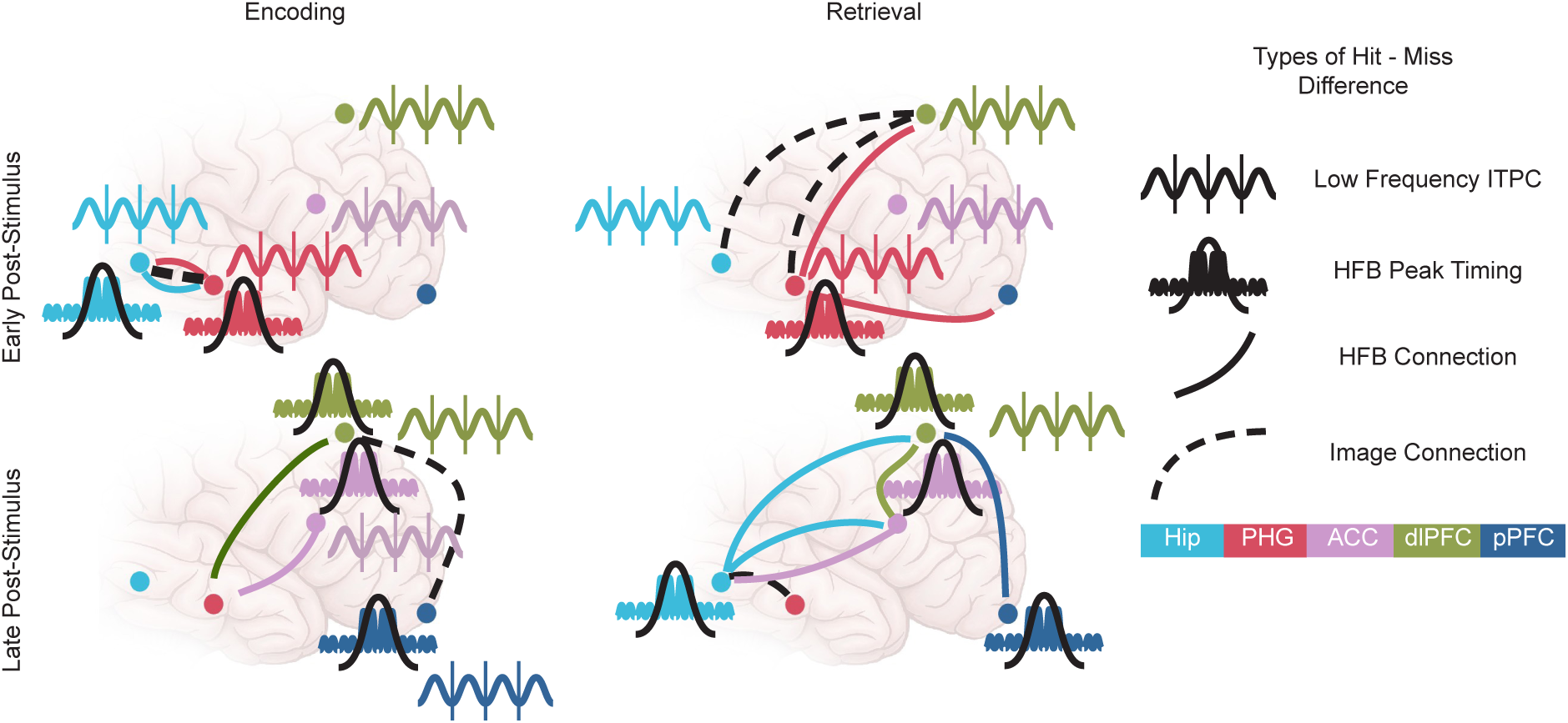
Schematic representation of key findings across all analyses.

Mean HFB peak latencies occurred late after stimulus onset in all regions of the PFC (Figure 7; left side, bottom panel) with the pPFC exhibiting the latest HFB peak. Locked to these HFB peaks, there were several signal components that increased during successful encoding. These included theta phase clustering across all three regions, and connectivity between the dlPFC and PHG and between the ACC and PHG for subsequently remembered images. Taken together, these physiological effects may represent inhibition of the PHG by the PFC in order to protect the Hip from interference. Indeed, the PHG is known to act as an informational gate to the Hip, and the anatomical connectivity of the PFC to the PHG could facilitate its control over such a gating mechanism ^53–57^. What’s more, at the time of the dlPFC’s HFB peak, low-frequency phase clustering and connectivity with the PHG both predicted individual memory performance (Figure 6), emphasizing the importance of these late effects for memory formation.

During successful retrieval (Figure 7; right side), PFC regions were active earlier in the trial, as has been reported previously ^7^, and connectivity was increased relative to encoding, as has been reported in rodents ^58^. Prior to image onset, successful retrieval was characterized by increased HFB and theta power in the dlPFC (Supplemental Figures 1 and 3). These mnemonic effects are not markedly different from those observed prior to image onset during encoding. These effects may index more domain-general processes like attentional or perceptual readiness rather than mnemonic processing itself ^34^.

Immediately after image onset (Figure 7; right side; top panel), hit trials were characterized by a complex flurry of differences from miss trials that contrasted sharply with the equivalent period during encoding when effects had been limited to the MTL. Increased connectivity was observed between the dlPFC and both MTL ROIs between -50ms and 200ms relative to image onset and concentrated between 2 and 3.5 Hz (Supplemental Figure 8). This time period and frequency range overlapped with retrieval-associated phase clustering in both MTL ROIs and the dlPFC, and these effects predicted individual memory performance (Figure 6; although dlPFC-Hip connectivity only yielded marginal significance r=.74; p=.06; n=7). Thus, whereas successful encoding was initiated by Hip theta signals, successful retrieval was initiated by an MTL-PFC network without a clear initiating node.

Later, after image onset, successful retrieval was characterized by near simultaneous HFB peak latencies in the Hip, ACC, dlPFC, and pPFC, which was in contrast to the temporal separation of these events observed during encoding. There were other notable differences in this time period from encoding. Whereas MTL-PFC connectivity during successful encoding was observed locked to the HFB peaks of the ACC and dlPFC and connected with the PHG, during successful retrieval, these connections were made with the Hip and were locked to the Hip’s HFB peak. It may be that these connections underlie the active processing of memory retrieval itself and the transfer of this information from the Hip to the PFC. However, these connections did not correlate with individual memory performance (ps>.5). It could be that late effects reflect recollection and earlier effects reflect familiarity. It is known that the Hip participates in both of these processes ^16,59,60^, but that they have different behavioral and event-related potential time courses ^61,62^. Specifically, familiarity is a feeling of knowing that a stimulus is old (e.g. recognizing acquaintance from stranger), and recollection is the additional retrieval of source information (e.g. hearing the acquaintance’s name) ^62,63^. Our recognition memory task did not require recollection per se, which may explain why effects observed during this stage of processing did not correlate with individual memory performance.

Across encoding and retrieval, the present results also recapitulated the observation that low-frequency phase clustering facilitates subsequent behavior through attentional orientation in declarative memory and other cognitive tasks ^64–68^. Extending this observation, PFC phase clustering was stronger in association with HFB peaks than with image onset, suggesting attentional orientation to internal information during mnemonic processing.

Our connectivity results both replicate and challenge prior reports. Of 98 significant differences in connectivity between ROIs as a function of memory outcome, 69 were such that connectivity was greater on hit trials, echoing previous reports that theta connectivity is elevated during successful memory ^31,49^. Did connectivity increase throughout the whole brain for successful encoding during these moments? We employed graph theoretic analysis to interrogate the overall network state during the limited time-frequency moments when connections between ROIs differentiated hit from miss trials. To our surprise, and contrary to prior results based on long temporal epochs, the overall network state was sparse at the precise moments when ROI-ROI connections were increased (Figure 5). This pattern was absent when connectivity was averaged across long temporal windows aligned to image onset. These results suggest that successful mnemonic processing at both encoding and retrieval is supported by a series of short-lived, sparsely connected network states. Similarly, it has been shown that decreased between-network communication is associated with better behavioral performance ^69^. This may indicate the ability to handle simple tasks, like the one studied here, with a modular (as opposed to integrated) network architecture ^70^.

Aligning analyses to HFB peaks is not dissimilar from the growing literature examining the role of “ripples” in cognitive processing ^27,71–73^. We do not claim here to be examining ripples since we did not have a minimum threshold for HFB power, a requirement for band limitation, a minimum cycle count, simultaneous sharp wave evaluation, or fine-grained anatomy ^74^. Indeed, examples of the individual HFB peak events analyzed here do not appear visually similar to examples of ripples reported in the literature. Nevertheless, averaging raw data aligned to the HFB peaks did yield grand average waveforms that appeared similar to sharp-wave ripple events (Supplemental Figure 11). More work is needed to understand the difference between transient increases in HFB power and ripple events. However, similar to the results reported in association with ripples, our findings emphasize that transient HFB signals are but one part of complex, multi-component events.

The present results should be interpreted with caution. First, it is likely that aspects of the brain dynamics reported here develop with age, or may vary with sex or other demographic characteristics. We chose to focus on an adolescent and early adult sample to balance maximization of sample size with minimization of developmental effects, yielding a tighter age distribution than many iEEG studies with similar n. Future research with larger sample sizes should examine how the mnemonic effects reported here may vary across the lifespan or as a function of sex or other demographic characteristics ^75–77^. Second, we chose to focus on a set of brain regions that are known to be involved in memory processing. However, other regions certainly participate as well, and future research should examine how the brain regions examined here interact with others. Third, we did not analyze phase-amplitude coupling (PAC). Given the focus on discrete dynamics, we believe that focusing on phase clustering at particular time points of high HFB power captures the relationship between HFB and theta signaling effectively. PAC analyses either require long temporal windows encompassing multiple cycles of the low-frequency signal or, if done at the singular time point of HFB peak across trials would be equivalent to HFB-locked phase clustering presented here. Finally, with any iEEG study there is a balance between anatomical specificity and sample size. In our analysis we focused on relatively large ROIs, and our analysis was agnostic to hemisphere. Anatomical variability at finer scales and between hemispheres certainly exists and likely introduced extra variability into our analysis. Fine anatomical differences would have introduced variance and decreased our chances of observing statistical significance. Future studies will likely reveal further nuance with sample sizes sufficient to consider finer anatomical distinctions.

In conclusion, we have found evidence for a sequence of transient processing states supporting recognition memory. These states are characterized by patterns of theta connectivity, phase resetting, and power, along with HFB activity changes, and they can be mapped meaningfully onto well-studied constructs from cognitive psychology. Intriguingly, many aspects of the sequence were detected with HFB-locked analyses, emphasizing the ephemeral and internally organized nature of mnemonic processing states. Lastly, although connections between key memory regions were strengthened during processing states, global connectivity was reduced, an effect that was lost when memory failed, suggesting a high degree of specificity in the effects reported here.

## STAR METHODS

### Experimental Model and Study Participant Details

Participants were 36 adolescents and adults (22 males; 13-28 years of age; M ± SD, 19.0 ± 4.7 years) undergoing intracranial monitoring as part of clinical management of seizures. We did not have access to participants’ ancestry, race, ethnicity, or socioeconomic status. Demographic and behavioral data are provided in Table 1. Subjects were selected from a larger pool based on non-pathologic sampling of regions of interest (ROIs; i.e. electrodes localized to the Hip, PHG, ACC, dlPFC, and/or pPFC and outside seizure onset zones ^80^), age greater than 13 years, and memory accuracy above chance (see below). There is partial overlap in subjects between this study and earlier studies using the same memory task ^32–34^. Subjects were recruited from Northwestern Memorial Hospital, the Ann & Robert H. Lurie Children’s Hospital of Chicago, the Children’s Hospital of Michigan, the University of California, San Diego Rady Children’s Hospital, the University of California, Irvine Medical Center, the University of California, Davis Medical Center, Nationwide Children’s Hospital, California Pacific Medical Center, and St. Louis Children’s Hospital. Written informed consent was obtained from subjects aged 18 years and older and from the guardians of all subjects younger than 18 years; written assent was obtained from subjects aged 13-17 years. All consent procedures were performed in accordance with the Declaration of Helsinki as part of the research protocol approved by the Institutional Review Board at each hospital.

### Method Details

#### Behavioral Task

Subjects performed a scene recognition memory task (Figure 1A) that has been used extensively to delineate the functional architecture of memory development with fMRI and electrocorticography (ECoG) ^32–34,75,81–85^. Subjects studied sets of 40 indoor and outdoor scenes, each shown for 3 s following a 0.5-s fixation interval. Stimuli were full-color pictures of natural scenes, half of high complexity and half of low complexity, characterized based on the number of object categories (over/under four) depicted ^33,86^. During the encoding phase, subjects were instructed to indicate verbally whether each studied item depicted an indoor or an outdoor scene. Responses were coded as correct or incorrect via offline review of individual audio recordings. A fixation cross remained on screen until a response was provided if none was provided during the 3-s scene presentation epoch. Per-trial RTs were automatically calculated by subtracting scene onset times from verbal response onset times. Analysis of electrophysiological data was restricted to trials in which scenes were correctly classified as indoor/outdoor, indicating the scenes were properly attended during the study block ^32^ ^33,34^. In addition, only trials with RTs below 3 seconds were considered.

The memory recognition test included all 40 scenes presented during the encoding block, intermixed in a randomized order with 20 new scenes. Following a 0.5-s fixation pretrial interval, each scene remained on screen until a response was given. Subjects verbalized an old/new judgment for each scene, which was coded as a hit (correct old), miss (new response to an old scene), correct rejection (CR; correct new), or false alarm (FA; old response to a new scene) via offline review of individual audio recordings. RTs were automatically calculated by subtracting scene onset times from verbal response onset times, and trials were excluded if no response was given.

This procedure was repeated twice, yielding total trial counts of 80 encoding trials, 80 retrieval trials with old images, and 40 retrieval trials with new images. All subjects completed a short practice run and at least one full encoding-test run. Recognition accuracy was calculated as hit rate minus false alarm rate ^32–34^. Only subjects with recognition accuracy greater than 0 were included in the present analysis.

The present analysis focuses on subsequent hit, subsequent miss, hit, and miss trials.

#### Electrode placement and localization

Macro-electrodes were surgically implanted for extra-operative recording based solely on the clinical needs of each patient. Electrodes were placed subdurally in 4- to 8-channel strips and 2-8 by 8 channel grids with 10-mm spacing (i.e., ECoG) or stereotactically in 8- to 18-channel tracks with 5- or 10-mm spacing (i.e., sEEG). Anatomical locations were determined by co-registering post-implantation computed tomography (CT) coordinates to pre-operative magnetic resonance (MR) images, as implemented in FieldTrip ^87^. Electrodes were localized in native space based on visual inspection of individual anatomy. For group-level visualization, electrode locations were transformed into standard MNI space.

#### Data acquisition and preprocessing

EEG data were acquired using Nihon Kohden Systems sampled at between .5 and 5 kHz. The BCI2000 software was used to acquire and store data in a subset of subjects. Raw EEG data were filtered with 0.1-Hz high-pass and 300-Hz low-pass finite impulse response filters, and 60-Hz line noise harmonics were removed using discrete Fourier transform. Continuous data were demeaned, epoched into 4.5-s trials (−1 to +3.5 s from scene onset), and manually inspected blind to electrode locations and experimental task parameters. Electrodes overlying seizure onset zones ^80^ and electrodes and epochs displaying epileptiform activity or artifactual signal (from poor contact, machine noise, etc.) were excluded, ensuring that data analyzed would represent healthy tissue ^88^. Neighboring artifact-free electrodes within the same anatomical structure were then bipolar referenced using consistent conventions (ECoG: anterior – posterior, sEEG: deep – surface) to form virtual channels anatomically located halfway between the two contributing electrodes ^39,89,90^. An electrode was discarded if it did not have an adjacent neighbor, its neighbor was in a different anatomical structure, or both it and its neighbor were in white matter. Bipolar referencing was used to minimize contamination from volume conduction ^91^. Data were then manually re-inspected to reject any trials with remaining noise. Functions from the FieldTrip toolbox for MATLAB were used for preprocessing routines ^92^.

#### Electrode selection by regions of interest

Bipolar channels were selected for analysis in the present study if they fell into one of five ROIs: pPFC, dlPFC, ACC, PHG, or Hip. The pPFC was defined as Broadman’s area 10. The dlPFC was defined as the posterior two-thirds of the superior and lateral surface areas of the superior frontal gyrus, the posterior two-thirds of the middle frontal gyrus. And the posterior one third of the inferior frontal gyrus. The ACC was defined as Broadman’s areas 32, 33, and 25, as well as the anterior half of area 24. The PHG was defined as the combined perirhinal, entorhinal, and parahippocampal cortices ^93^. The Hip was defined as the entire volume of the dentate gyrus, fields of cornu ammonis, and subiculum. Across subjects, there were 35 channels in the pPFC, 247 in the dlPFC, 44 in the ACC, 79 in the PHG, and 40 in the Hip.

### Quantification and Statistical Analysis

#### Signal processing analysis approach

Although the analysis presented here focuses on five ROIs, key signal components were extracted for all recorded channels. This approach facilitated graph theoretical analysis (described below), but was computationally intensive, requiring parallelization of analysis code for computation on a high-power cluster (HPC). Coding principles to facilitate this analysis approach have been described previously ^94^. Briefly, each participant’s data were divided into a set of individual channel files. Signal processing was then performed on these individual channel files in parallel, and inferential statistics were performed on ROI groups of channel files after signal processing was complete. Key signal processing steps described in the remainder of this sub-section were all executed in the function singleChanPipeline. All signal components were down-sampled after extraction to a sampling rate of 40 Hz to reduce file sizes and computation time.

High-frequency broadband (HFB) time series were extracted. Data were narrowband filtered using the Fieldtrip ^92^ function ft_specest_mtmconvol between 70 and 150 Hz using ‘dpss’ (i.e., multitaper analysis ^95^) for the taper argument and padding equal to the next power of two divided by the sample rate. Filtered data were converted into power time series by absolute valuing and squaring. The mean was taken across tapers. Power time series were z-scored within frequency based on the mean and standard deviation of a pre-stimulus baseline period of -450 to -50 ms relative to image onset using a bootstrap procedure implemented in myChanZscore and described previously ^33,34,37,39,89,90^. Finally, a single HFB time series was obtained by taking the mean across frequency of the z-scored power time series. See functions getHFB, myChanZscore, and getChanMultiTF for more detail.

Only channels whose HFB time series exhibited evidence of task reactivity were included in inferential statistics and plots. Each channel was deemed reactive if its mean HFB time series across trials exceeded z=1.96 for any continuous 50 ms period between -50 ms and 2000 ms relative to image onset or -1500 ms and 500 ms relative to behavioral response ^96^. Reactivity was assessed at both encoding and retrieval. Channels that were reactive for any one period were used in all analyses. Reactivity was calculated with data aggregated across all trial types. See functions checkForThreshold_100 and reactiveTest_100 for details.

The latency of peak HFB activity was calculated for each trial. Each trial was smoothed with a 275 ms Gaussian kernel convolved with the HFB time series. The latency of the peak of the smoothed data between image onset and the behavioral response was extracted. See function gausLat for details.

Morlet Wavelet convolution in the frequency domain was used to extract narrowband complex time series at 100 logarithmically spaced frequencies between 2 and 80 with corresponding standard deviations ranging from 3 to 10 ^97^. For each frequency, the resulting analytic signal was converted into a phase time series using the MATLAB function angle and a power time series using absolute value and squaring. Power time series were z-scored within frequency in the same way as HFB time series. See functions getChanTrialTF and myChanZscore for details. Inter-trial phase clustering (ITPC) was calculated using the phase time series at the time of inferential statistics ^97^.

Pairwise phase consistency (PPC) ^40^ was extracted between all pairs of channels for all trial types across 20 logarithmically spaced frequencies between 2 and 25 Hz. These connectivity metrics were calculated across trials for each time point, with trials aligned in time to the onset of the image. In addition, these metrics were also calculated across trials for each time point with trials aligned to the HFB peak latency. Because HFB latency was different for different channels, the HFB latency-aligned connectivity analysis was non-symmetric between regions. For example, connectivity was calculated between the Hip and PHG using time aligned to the peak of the HFB in the Hip, and it was also calculated using time aligned to the peak of the HFB in the PHG. Calculation of PPC was done using custom code. Different trial types were analyzed separately. See function getChanISPC for details.

#### HFB-locked and image-locked temporal alignment

Inferential statistics were done in two ways for all signal components. HFB-locked analyses used the latency of the HFB peak in each trial as t=0 and data points surrounding the HFB peak were extracted for analysis +/-500ms around this time point for each trial. Image-locked analyses used the latency of the image onset on the computer screen as t=0 and data points surrounding the image onset were extracted for analysis -450 to 3000 ms for each trial.

#### Within-ROI inferential statistics

Within-ROI hit/miss differences were examined independently for each ROI and for encoding/retrieval using a standardized method across all signal components and both HFB-locked and image-locked temporal alignments. Specifically, this method was used for HFB power, low-frequency power, low-frequency phase reset (ITPC), and PPC data. Inferential analysis focused on measuring differences between (subsequent) hit vs. miss trials. Linear mixed-effects (LME) models were applied independently to model different signal components and for encoding and retrieval data. For all analyses hit/miss was a fixed effect and channel and subject ID were random effects ^98,99^.

Because signal components varied across both time and frequency, LME models were applied at every time and frequency point independently. To correct for multiple comparisons, cluster-based permutation testing was applied ^100,101^. Specifically, the channel mean for (subsequent) hit and miss trials was calculated for each measure at each time-frequency point. Next, the hit and miss identity of each channel’s observed values was shuffled within channel and the model was refit. This process was repeated 2,500 times. The function cluster_test was used to evaluate significance of the observed model output versus the permutation outputs ^100^. Specific functions to generate the permutations and fit the LME models were written for each signal component separately: HFBsingleTrialPipeline.m, connectivityPipeline.m, TFphaseTrialpipeline.m, TFsingleTrialpipeline.m.

#### HFB latency between ROI inferential statistics

HFB latency values for all individual trials along with subject ID, channel ID, encoding/retrieval, hit/miss, and behavioral RT were aggregated and exported from MATLAB for modeling in R using the Tidyverse for general data handling ^102^, and the lme4 package for modeling ^103^. HFB latency was modeled as a function of the fixed effects of hit/miss * encoding/retrieval * region with subject ID and channel ID used as random effects (Figure 2D). Holm-corrected pairwise contrasts were evaluated to aid interpretation of significant effects in the model using the R function emmeans ^104^.

A major challenge with this analysis was that different participants contributed channels from different ROIs. Thus, it was difficult to know if differences in the relative timing of HFB latency between ROIs were due to differences in regional brain function or differences between individual subjects. To mitigate this confound, this analysis was carried out in 3 ways. First, a bias-corrected measure of HFB latency was modeled (main text). Specifically, HFB latency was divided by RT to provide a measure that ranged from 0 to 1 and accounted for individual differences in RT. Second, the raw HFB latency was modeled (Supplemental Figure 2A). Third, a subset of trials with matched RTs were selected and these were used for modeling (Supplemental Figure 2B). Fourth, in the matched subset of trials HFB latency was divided by RT and these bias-corrected values of the matched trials were used for modeling (Supplemental Figure 2C). The Anova function in R was used to extract chi-squared and p values from these models using Type II Wald tests. The emmeans function with holm adjustment for multiple comparisons was used to perform pairwise contrasts between regions within different trial types and also between different trial types within region.

As an additional check, for pairwise sets of ROIs, the single-trial HFB latencies were compared in simultaneously recorded pairs of channels (Supplemental Figure 2D). Histograms were plotted of the relative HFB latency for these within-subject comparisons. In addition, ROI label was shuffled to generate a null distribution. The difference between these distributions was evaluated with KS-test. A key question was whether the directionality of these within-subject comparisons would corroborate the results of the omnibus between-subject analyses described in the previous paragraph. All analyses in this section were executed in Latency_LME_modeling.Rmd.

#### HFB-locked power spectrum between ROI inferential statistics

Visual inspection of time-frequency plots centered on HFB peaks indicated that power increases were highly concentrated at the time point of the peak itself (Supplemental Figure 3A and B), and so instantaneous estimates of the power spectra at this time point were examined alone (Supplemental Figure 3C). Differences in the power spectra observed were explored using the lmer function in R. Power was modeled as a function of frequency (2-80 Hz; modeled with 8 splines) * region * encoding/retrieval with subject ID and channel ID used as random effects. The Anova function in R was used to extract chi-squared and p values from these models using Type II Wald tests. These analyses are implemented in GAM_analysis_HFB_spectra.Rmd.

Because frequency was a continuous variable and was modeled non-linearly, it was not straightforward to extract post-hoc contrasts to explain significant effects in the full model. Instead, we took a descriptive approach. We reasoned that if peak frequencies in the power spectra varied between ROIs and conditions, then these differences in power likely contributed to the effects in the full model. To do this, a low spectral peak was identified for each ROI during encoding and retrieval by identifying the frequency at which power was maximal between 2 and 20 Hz. A medium spectral peak was identified by searching for a second peak above the low peak and below 34 Hz. The presence of a medium spectral peak was defined to exist when the maximum power value higher than the first trough after the low peak and below 34 Hz was a higher value than the power at either of the endpoints of this range. The key idea is that any differences in the shape of the power spectra will drive differences in the omnibus model. Thus, the particular choice of descriptive methodology does not influence whether or not power values varied as a function of frequency, region, or encoding/retrieval. The point is simply to describe how the shape of the power spectrum changed across conditions.

#### Contrasting ITPC between image and HFB locked analyses (IMG-HFB index)

An IMG-HFB index was used to quantify the relative strength of ITPC differences in HFB-locked versus image-locked analyses (Figure 3C-E). To do this, for each region a frequency of maximal ITPC was determined using data from hit trials with HFB-locked time alignment. The Raleigh’s Z magnitude of ITPC at this target frequency in HFB-locked data for hit and miss trials was extracted for each channel. For image-locked data, ITPC values were extracted for each channel at the same target frequency for hit and miss trials and at the time point of maximal ITPC in each trial type. This resulted in one ITPC value for each of four conditions for every channel: HFB-locked hits, HFB-locked misses, image-locked hits, and image-locked misses. Corresponding HFB and image-locked values were used to create an index score for each channel and trial type reflecting the relative strength of ITPC values across the two time alignment methods:

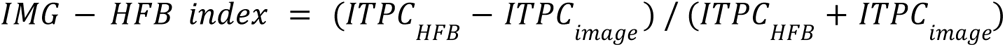

This index could take on values between -1 and +1 where -1 indicated that image-locked ITPC was much stronger than HFB-locked ITPC.

IMG-HFB index values were modeled using linear mixed effects modeling as a function of region, hit/miss, and encoding/retrieval. This analysis followed the same approach as the latency analysis described above and was executed in HFB_IMG_index.Rmd.

#### Visualization of connectivity results

To aid interpretation, ROIs were plotted schematically as dots, and connections that significantly increased in strength during hit trials over miss trials were plotted as lines connecting ROIs (Figure 4E and F). Heatmaps displaying granular time-frequency plots of PPC for individual connections are available in Supplemental Figures 6-9; see supplemental Figure 5 for equivalent figures displaying significant negative connectivity differences). These schematic connectivity plots were generated for HFB-locked (Figure 4E) and image-locked (Figure 4F) analyses separately, and connections were further grouped by time periods and encoding versus retrieval.

#### Graph theoretic analysis

Every significant difference between two ROIs in PPC between hit and miss trials was termed a mnemonic connection. The mean time and frequency of the significant difference was extracted. Using this time and frequency point, the full channel X channel connectivity matrix was extracted for every participant with channels in both ROIs of the mnemonic connection under consideration. These full channel X channel matrices included many channels outside ROIs, but the goal of this analysis was to understand the whole-brain network state at the moments of mnemonic connections. Channel X channel matrices were extracted for hit and miss trials independently. Mnemonic connections were assessed for encoding and retrieval separately.

Next, three graph theoretic measures were calculated for each channel X channel matrix using functions from the Brain Connectivity Toolbox ^42^. Weighted graph analysis treats each channel as a node and PPC values as the strength of the connecting edges of the network. Unweighted graph analysis is similar but requires that connections be binarized into values of either 0, indicating no connection along a possible edge, or 1, indicating a connection along a possible edge. When necessary, channel X channel connectivity matrices were binarized with a threshold of PPC > .1 => 1; PPC < .1 => 0, but the choice of threshold did not qualitatively affect results.

Analysis focused on three measures. The weighted characteristic path length was calculated using the BCT functions charpath and distance_wei and described the average distance of paths between any two nodes in the network. The weighted strength was calculated using the BCT function strengths_und and was simply the sum of PPC values observed for each channel. The unweighted strength was calculated as the average proportion of edges with PPC > .1 for each channel. Strength values were averaged over channels to obtain a single estimate for the full network.

Although channel coverage varied over participants, it is important to note that our main findings from this analysis focused on hit/miss differences rather than on the absolute value of graph measures (Figure 5). The LME approach models independent random intercepts for each participant, which accounts for the biasing difference of channel coverage, allowing for valid inferences about fixed effects of hit/miss. By contrast, regional differences in these measures must be interpreted with caution as different participants contributed observations from different regions. As such, regional differences are presented in the supplement only (Supplemental Figure 10).

LME modeling of graph measures followed a similar approach as presented above for HFB latency and each graph measure was modeled as a function of region * hit/miss * encode/retrieve in graphMeasures.Rmd.

These were weighted characteristic path length, unweighted strength, and weighted strength. Weighted characteristic path length uses the PPC values as edge strengths signifying

These channel X channel connectivity matrices were extracted and graph measures combined for statistical analysis in connectionGraph.m and finalGraphDatCombine.m. LME modeling was done in graphMeasures.Rmd.

#### Correlations with memory performance

For every significant difference between hit and miss trials across all signal components, individual memory performance (measured using d’ ^105^) was compared with individual differences between hit and miss trials. To do this, the significant cluster of time and frequency points for each hit miss difference was used as a mask to select the relevant time-frequency extent of the difference. For each participant, this mask was used to select data points from the mean hit signal and from the mean miss signal. Next, a difference score between hit and miss trials was calculated across the selected data points and the mean difference was calculated. When a participant had more than one channel that contributed to a significant difference, the mean hit/miss difference was calculated across channels for that participant. In this way, each participant contributed a single memory performance value and a single hit/miss signal difference value. Because of differences in channel coverage, the n associated with each signal component and regional difference varied. These values were extracted throughout publicationFigs.m and exported for analysis in R in the script allSig.Rmd.

Next, individual hit/miss signal differences and memory performance values were correlated with each other. For all significant correlations (p<.05), a leave one out procedure was performed. For this procedure, each participant was left out one at a time and the correlation was recalculated. Only those correlations that were significant in the full group at the .05 level and that maintained significance at the .10 level after removal of any one participant are reported in Figure 6. Given that n was often small and that correlation values can vary dramatically with single data points when n is small, this procedure ensured that none of the reported correlations were reliant on a single participant.

This analysis was exploratory and its goal was not to establish causal relationships between physiological measures and memory performance. Rather, the goal of this analysis was to aid interpretation by providing an additional statistical check beyond the rigorous LME and cluster permutation testing to which each hit/miss difference had already been subjected. In this way, from the large number of significant effects discovered and reported here, it was possible to highlight significant differences of particular promise for interpretation and future investigation.

---

**Supplemental Figure 1.** HFB power changes aligned to image onset and HFB peak. **A**. Each line plot displays the mean time series of the HFB response across channels within different regions. For all panels, the x-axis displays time relative to image onset, and the y-axis displays HFB power in z-scored units. Encoding and retrieval data are plotted along the top and bottom rows, respectively. Orange and blue lines are the average time series of (subsequent) hit and miss trials, respectively. vertical gray shaded regions indicate p<.05 for the difference between hit and miss after cluster correction. Successful encoding was associated with elevated HFB activity prior to image presentation in the dlPFC, and ∼500 ms after image presentation and after the indoor/outdoor behavioral response in the pPFC. Successful retrieval was associated with elevated HFB activity prior to image presentation in the dlPFC, but lower HFB activity for failed memory late in the trial in both the dlPFC and pPFC. The PHG exhibited effects during both encoding and retrieval, but these were much later and smaller in magnitude than the visual response. Colored shaded regions indicate the standard error of the mean. Colored vertical dashed lines indicate mean reaction times for hit trials (blue) and miss trials (orange). **B**. Each grouped scatter plot displays the mean power of HFB peak for each channel grouped by region. Mean z-scored HFB power at the time of the HFB peak is displayed on the y axis. Error bars display the 83% confidence interval around model estimates ^78,79^. Linear mixed effects modeling of these data revealed main effects of encode/retrieve (χ^2^(1) = 188, *p* < 2*e* − 16) and hit/miss (χ^2^(1) = 11, *p* =. 0008), and all four interaction terms: hit/miss by encode/retrieve (χ^2^(1) = 6, =. 016), hit/miss by region (χ^2^(1) = 11, *p* =. 02), encode/retrieve by region (χ^2^(1) = 14, *p* =. 007), and hit/miss by encode/retrieve by region (χ^2^(4) = 20, *p* =. 0005). Holm-corrected pairwise contrasts revealed that in the PHG, Hip, dlPFC, and pPFC, the peak HFB power was higher during retrieval than encoding. Mnemonic effects were evident in the ACC and PHG, with higher peak power during failed encoding than successful encoding. **C**. Mean HFB power time series using the same conventions as in A except with time centered around the latency of the HFB peak. Both panels display data from the encoding phase of the experiment, with ACC data displayed on top and PHG data below. Note that although the image-locked mean HFB activity in the ACC appeared flat (panel A), its peak activity was no smaller than in any other region.

**Supplemental Figure 2.** Timing of HFB peak latency can be measured in several ways, but all reinforce the same general interpretation. **A.** Each grouped scatter plot displays the mean time of peak HFB latency for each channel grouped by region. Time relative to image onset is displayed in milliseconds on the y axis. Error bars display the 83% confidence interval around model estimates ^78,79^. **B.** Similar to A except a subset of trials was selected such that reaction time was matched at the individual trial level. **C.** Similar to B, except that a subset of trials was selected, such that reaction time was matched at the individual trial level, and normalized time was used. **D**. The asynchrony of HFB latency within trial is displayed for pairs of channels recorded simultaneously within individual participants. The four groups of figures represent the behavioral conditions indicated by the large font labels in the left and top margins. The x axis shows the difference in timing of the HFB peak observed on single trials at pairs of simultaneously recorded channels. Positive values correspond to the brain region indicated at the top of the column having had an earlier HFB peak latency. Negative values correspond to the brain region indicated at the side of the column having had an earlier HFB peak latency. The y axis shows the proportion of trials observed to have a given peak latency asynchrony. The color of the line represents the brain region with the earlier latency (as judged by proportion of trials). The dashed line shows the latency asynchronies observed when region identity is shuffled prior to calculating latency asynchrony. The percent of trials where the row region was the leader is indicated in the upper left of each plot. The percent of trials where the column region was the leader is indicated in the upper right. Note, these values do not sum to 100% because ties were discounted. The pairwise contrasts observed here recapitulate the order observed in Figure 2D of the main text.

**Supplemental Figure 3.** TF power differences between hit and miss trials across encoding and retrieval. **A**. heatmaps display frequency in Hertz on the y axis and time in milliseconds relative to HFB peak on the x axis. Color indicates power z-scored within frequency. The top row reflects data collected during subsequent hit trials. The bottom row reflects data collected during subsequent miss trials. **B**. Similar to A except for retrieval. **C**. To facilitate comparison between hit and miss trials, line plots display power spectra at the time point of the HFB peak. For all panels, the x-axis displays frequency, and the y-axis displays power in z-scored units. Encoding and retrieval data are plotted along the top and bottom rows, respectively. Orange and blue lines are the average spectra of (subsequent) hit and miss trials, respectively. vertical gray shaded regions indicate p<.05 for the difference between hit and miss spectra after cluster correction. Colored shaded regions indicate the standard error of the mean. Successful encoding elicited greater power in the Hip between 2.0 and 2.9 Hz. Failed retrieval elicited greater power in the PHG between 4.4 Hz and 15 Hz. Although other ROIs did not exhibit hit/miss differences, power spectra nevertheless appeared different across regions and between encoding and retrieval. These differences were explored using linear mixed effects modeling of power as a function of frequency (2-80 Hz; modeled with 8 splines), region, and encoding/retrieval. All interactions between fixed effects were significant (χ^2^ (4 − 32) > 66, *maximum p* < 2*e* − 10). Several factors may explain these interactions. First, an interaction between frequency and region may have been driven by regional variations in the frequency of maximum power in the lower range (2-20 Hz; Table 1). Second, an interaction between encode/retrieve and region may have been driven by higher power in the Hip during retrieval than during encoding. Third, a three-way interaction may have been driven by secondary power increases between 24 and 32 Hz in the Hip, ACC, and dlPFC during retrieval (Table 1). These results emphasize that HFB peaks constituted physiological events, and not statistically extreme values. **D**. Similar to A, except time on the x axis is represented in seconds relative to the image onset. **E**. Similar to D, except for retrieval. **F**. Heatmaps display the mean difference in z-scored power between hit and miss trials. For all panels, the x-axis displays time relative to image onset, and the y-axis displays frequency. White outlined areas indicate p<.05 for the difference between hit and miss power after cluster correction. In the Hip and dlPFC, there were positive subsequent memory effects in 2 to 10 Hz power just before and during image onset (-450 to 100 ms). In the PHG, there was a negative memory effect in 2 to 10 Hz later after image onset at both encoding (550 to 1550 ms) and retrieval (925 to 1450 ms). These decreases partially overlapped with the visual response observed in the PHG’s HFB power (Supplemental Figure 1A). Finally, the pPFC exhibited a late (1225 to 2775 ms) positive subsequent memory effect during encoding across 2 to 10 Hz. At retrieval, the pPFC exhibited a much earlier (-100 to 600 ms) positive memory effect and a later (950 to 1975 ms) negative memory effect.

**Supplemental Figure 4.** Distributions of phase preference of phase clustering within channels. **A**. Histograms display the proportion of channels on the y axis and the phase of either the 3 Hz (left columns) or 6.5 Hz (right columns) component of the signal. For each channel, the mean circular phase was calculated across trials at the time point of the HFB peak for hit (blue) and miss (orange) trials separately. The superimposed sin wave on each plot indicates which phase values correspond to the peak, trough, or intermediate positions in the oscillation. The dashed-dotted lines indicate uniform distribution. The top row of each panel displays encoding data. The bottom row displays retrieval data. Each panel displays data for a different region. Notice that the Hip exhibited a preference for HFB activity to occur at the trough of the 3 Hz oscillation during successful memory trials at both encoding and retrieval, but the PHG exhibited a preference for the peak. **B**. Similar to A, but mean phase was measured at the time point of maximal ITPC relative to image onset. Notice that phase preferences evident in MTL regions in C appear weaker when measured relative to image onset.

**Supplemental Figure 5.** Negative mnemonic connections. Plots A and B correspond to main text Figure 4 E and F, respectively. However, while the schematics in Figure 4 E and F display only connections that exhibited increased strength for hit trials, this figure displays only connections that exhibited decreased connectivity strength for hit trials relative to miss trials. Note that far fewer of these negative connections were detected than the positive connections shown in Figure 4.

**Supplemental Figure 6.** Inter-regional connectivity changes associated with memory during encoding. Analysis aligned to image onset. In all panels, subplots have time relative to image onset (t=0) on the X-axis and frequency on the Y-axis. The color scale indicates the strength of connectivity between the regions indicated by the row and column of the subplot. **A.** Connectivity during subsequent hit trials using pairwise phase consistency. **B.** Connectivity during subsequent miss trials measured using pairwise phase consistency. **C.** The difference between hit and miss trials. Warm colors indicate stronger connectivity during hit trials. Color scale represents t-values. White outlines indicate cluster-corrected significant differences between hit and miss trials.

**Supplemental Figure 7.** Inter-regional connectivity changes associated with memory during encoding. Analysis aligned to HFB peak of the region indicated by the row position of each subplot. In all panels, subplots have time relative to HFB peak (t=0) on the X-axis and frequency on the Y-axis. The color scale indicates the strength of connectivity between the regions indicated by the row and column of the subplot. **A.** Connectivity during subsequent hit trials measured using pairwise phase consistency. **B.** Connectivity during subsequent miss trials measured using pairwise phase consistency. **C.** The difference between hit and miss trials. Warm colors indicate stronger connectivity during hit trials. Color scale represents t-values. White outlines indicate cluster-corrected significant differences between hit and miss trials.

**Supplemental Figure 8.** Inter-regional connectivity changes associated with memory during retrieval. Analysis aligned to image onset. In all panels, subplots have time relative to image onset (t=0) on the X-axis and frequency on the Y-axis. The color scale indicates the strength of connectivity between the regions indicated by the row and column of the subplot. **A.** Connectivity during hit trials measured using pairwise phase consistency. **B.** Connectivity during miss trials measured using pairwise phase consistency. **C.** The difference between hit and miss trials. Warm colors indicate stronger connectivity during hit trials. Color scale represents t-values. White outlines indicate cluster-corrected significant differences between hit and miss trials.

**Supplemental Figure 9.** Inter-regional connectivity changes associated with memory during retrieval. Analysis aligned to HFB peak of the region indicated by the row position of each subplot. In all panels, subplots have time relative to HFB peak (t=0) on the X-axis and frequency on the Y-axis. The color scale indicates the strength of connectivity between the regions indicated by the row and column of the subplot. **A.** Connectivity during hit trials measured using pairwise phase consistency. **B.** Connectivity during miss trials measured using pairwise phase consistency. **C.** The difference between hit and miss trials. Warm colors indicate stronger connectivity during hit trials. Color scale represents t-values. White outlines indicate cluster-corrected significant differences between hit and miss trials.

**Supplemental Figure 10.** Additional graph analysis results. **A.** Box plot display connectivity between the ACC and Hip for all channel pairs that spanned the two regions (open blue circles) and participants (red circles) averaged across the same time and frequency windows as in Figure 5A.i and A.ii. Note that although miss trials were characterized by more densely connected overall graphs (A.ii), connectivity between the ACC and Hip was weaker during miss trials for all participants. **B.** The histogram displays the strength of all connections for the representative participant shown in Figure 5A averaged across the long temporal epoch (corresponding to Figure A.iii and A.iv). Note that with lower variance in the hit distribution, a higher central tendency in the hit distribution, and similar representation at very strong connectivity strengths between hit and miss trials (see inset), a z-scored analysis would replicate prior findings of stronger connectivity for hit trials ^31^. Overall, for this example participant, 52% of all possible connections exhibited greater connectivity values for successful encoding over failed encoding, and the corresponding value was 54% for retrieval trails. It is interesting that overall connectivity was stronger during hit trials when examined over the longer temporal window. **C.** Same as in Figure 5B but with data sorted by whether significant connections were detected in HFB-aligned or image-aligned analyses. There was a modest effect such that graphs exhibited more connectivity for HFB-locked analyses than image-locked: shorter characteristic path length, increased weighted strength, and increased unweighted strength; χ^2^ (1) > 12, *maximum p* < .0005. **D.** Boxplots display results of graph analysis for each region and timeset combination separately during encoding. Top, middle, and bottom panels display characteristic path length, weighted node strength, and unweighted node strength respectively. The colors of dots represent the region of each connection partner. Asterisks indicate significant (p<.00a01) holm-corrected comparisons between HFB-aligned and image-aligned values within region. Notice that Hip and pPFC graphs are more connected when aligned to HFB peaks during successful encoding. This is true for the pPFC during failed encoding as well. **E.** Similar to D except for retrieval.

**Supplemental Figure 11.** Example HFB peaks do not look like sharp wave ripples, but their mean does. **A-D** Line plots display individual HFB peak events. The raw timeseries is displayed in black and the timeseries bandpass filtered to between 70 and 150 Hz is displayed in purple. Panel ii displays a zoomed in time scale relative to panel i. Time is displayed on the x axis relative to the time point of HFB peak detection. The y axis displays the electrical potential measured in microvolts. All randomly chosen events are from hippocampal channels. Each event is recorded in a different participant. **E.** The line plot displays the mean of the raw hippocampal encoding timeseries aligned to the nearest trough of the HFB signal relative to the peak of the HFB power for hit and miss trials separately. Notice that although no individual events bear visual resemblance to classical sharp wave ripples, these mean events do.

## Author Contributions

Conceptualization, AJOD, NO, ELJ; Methodology, AJOD, NO, ELJ; Software, AJOD, JPK; Formal Analysis, AJOD, ZRC; Investigation, AJOD, ZRC, SMG, QY, ELJ; Resources, EA, SUS, JMR, JYW, SKL, JSR, JJL, OKM, SS, SA, DK, PBW, KDL, PB, JLR, IS, FG; Data curation, SMG, QY, PV, NO, ELJ; Writing-original draft, AJOD; Writing-reviewing & editing, AJOD, JPK, ELJ; Visualization, AJOD; Supervision, RTK, NO, ELJ; Project Administration, ELJ; Funding acquisition, RTK, NO, ELJ

## Declaration of Generative AI and AI-assisted Technologies

chatGPT was used for first draft code writing in many instances. However, no code was used without careful evaluation and editing by the first author.

## Acknowledgements

We thank the participants who volunteered their time and all of the clinical staff who helped facilitate data collection. This research was supported in part by the computational resources and staff contributions provided for the Quest high-performance computing facility at Northwestern University, which is jointly supported by the Office of the Provost, the Office for Research, and Northwestern University Information Technology. REDCap is supported at FSM by the Northwestern University Clinical and Translational Science (NUCATS) Institute. Research reported in this publication was supported, in part, by the National Institute of Health’s National Center for Advancing Translational Sciences, Grant Number UL1TR001422. The content is solely the responsibility of the authors and does not necessarily represent the official views of the National Institutes of Health. This work was funded by grants from the National Institutes of Health (NINDS R00NS115918, NIMH R01MH107512, NINDS R01NS021135, NIBIB P41EB018783, NINDS T32NS047987).

## Resource Availability

### Data and Code Availability

Processed data with all necessary summary statistics for all inferential statistics and figure generation along with custom Matlab and R code will be posted after peer review.

