## supplemental figures for "Declarative Memory Through the Lens of Single-Trial Peaks in High-Frequency Power"

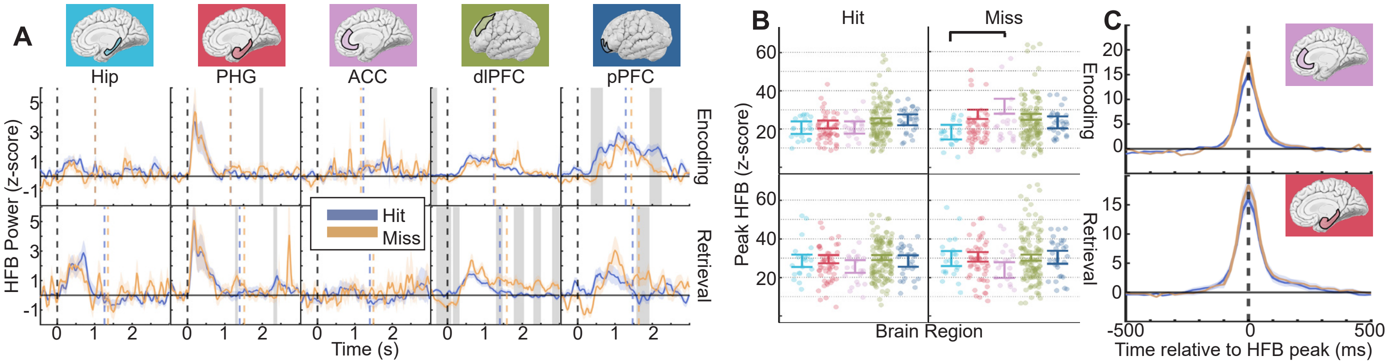


Supplemental Figure 1. HFB power changes aligned to image onset and HFB peak. A. Each line plot displays the mean time series of the HFB response across channels within different regions. For all panels, the x-axis displays time relative to image onset, and the y-axis displays HFB power in z-scored units. Encoding and retrieval data are plotted along the top and bottom rows, respectively. Orange and blue lines are the average time series of (subsequent) hit and miss trials, respectively. vertical gray shaded regions indicate p<.05 for the difference between hit and miss after cluster correction. Successful encoding was associated with elevated HFB activity prior to image presentation in the dlPFC, and ~500 ms after image presentation and after the indoor/outdoor behavioral response in the pPFC. Successful retrieval was associated with elevated HFB activity prior to image presentation in the dlPFC, but lower HFB activity for failed memory late in the trial in both the dlPFC and pPFC. The PHG exhibited effects during both encoding and retrieval, but these were much later and smaller in magnitude than the visual response. Colored shaded regions indicate the standard error of the mean. Colored vertical dashed lines indicate mean reaction times for hit trials (blue) and miss trials (orange). B. Similar to Figure 2D except displaying HFB power rather than latency. Error bar conventions are as in Figure 2D. Mean z-scored HFB power at the time of the HFB peak is displayed on the y axis. Linear mixed effects modeling of these data revealed main effects of encode/retrieve (2(1)=188, p<2e-16) and hit/miss (2(1)=11, p=.0008), and all four interaction terms: hit/miss by encode/retrieve (2(1)=6, =.016), hit/miss by region (2(1)=11, p=.02), encode/retrieve by region (2(1)=14, p=.007), and hit/miss by encode/retrieve by region (2(4)=20, p=.0005). Holm-corrected pairwise contrasts revealed that in the PHG, Hip, dlPFC, and pPFC, the peak HFB power was higher during retrieval than encoding. Mnemonic effects were evident in the ACC and PHG with higher peak power during failed encoding than successful encoding. C. Mean HFB power time series using the same conventions as in A except with time centered around the latency of the HFB peak. Both panels display data from the encoding phase of the experiment with ACC data displayed on top and PHG data below. Note that although the image-locked mean HFB activity in the ACC appeared flat (panel A), its peak activity was no smaller than in any other region.

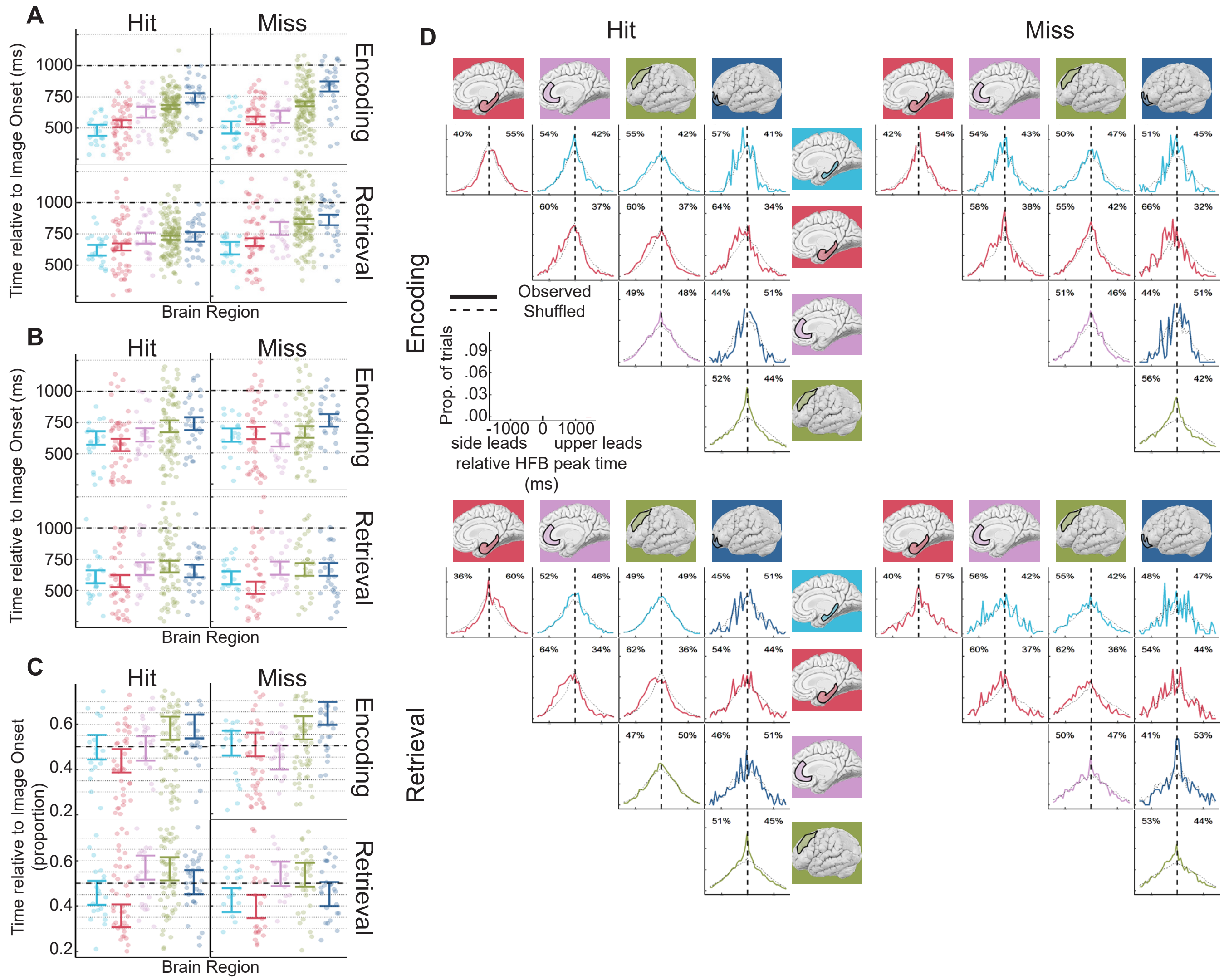


Supplemental Figure 2. Timing of HFB peak latency can be measured in several ways, but all reinforce the same general interpretation. A. Similar to Figure 2D except raw time was used. B. Similar to A except a subset of trials was selected such that reaction time was matched at the individual trial level. C. Similar to B except that a subset of trials was selected such that reaction time was matched at the individual trial level, and normalized time was used. D. The asynchrony of HFB latency within trial is displayed for pairs of channels recorded simultaneously within individual participants. The four groups of figures represent the behavioral conditions indicated by the large font labels in the left and top margins. The x axis shows the difference in timing of the HFB peak observed on single trials at pairs of simultaneously recorded channels. Positive values correspond to the brain region indicated at the top of the column having had an earlier HFB peak latency. Negative values correspond to the brain region indicated at the side of the column having had an earlier HFB peak latency. The y axis shows the proportion of trials observed to have a given peak latency asynchrony. The color of the line represents the brain region with the earlier latency (as judged by proportion of trials). The dashed line shows the latency asynchronies observed when region identity is shuffled prior to calculating latency asynchrony. The percent of trials where the row region was the leader is indicated in the upper left of each plot. The percent of trials where the column region was the leader is indicated in the upper right. Note, these values do not sum to 100% because ties were discounted. The pairwise contrasts observed here recapitulate the order observed in Figure 2D of the main text.

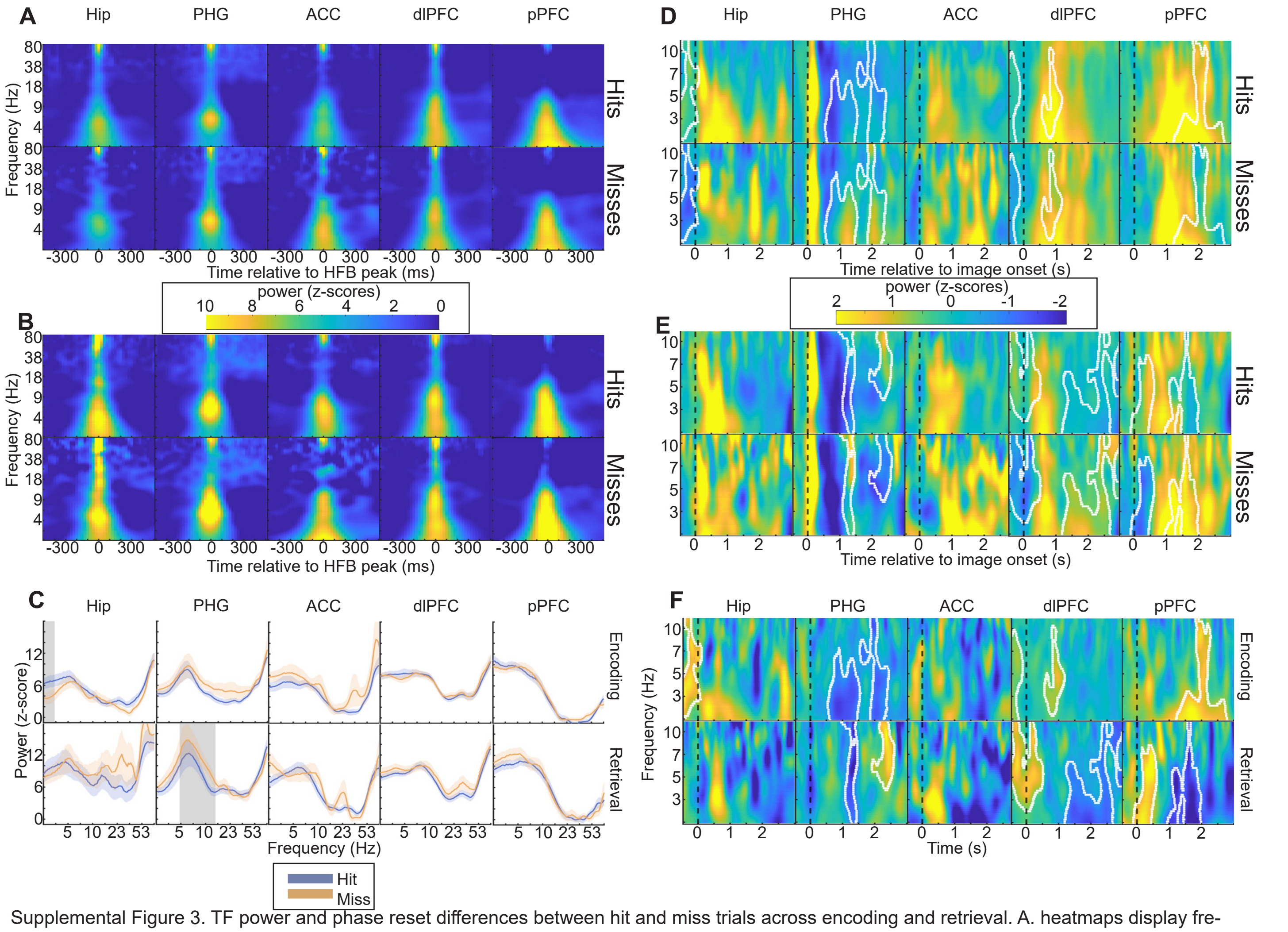


Supplemental Figure 3. TF power and phase reset differences between hit and miss trials across encoding and retrieval. A. heatmaps display frequency in Hertz on the y axis and time in milliseconds relative to HFB peak on the x axis. Color indicates power z-scored within frequency. The top row reflects data collected during subsequent hit trials. The bottom row reflects data collected during subsequent miss trials. B. Similar to A except for retrieval. C. To facilitate comparison between hit and miss trials, line plots display power spectra at the time point of the HFB peak. For all panels, the x-axis displays frequency, and the y-axis displays power in z-scored units. Encoding and retrieval data are plotted along the top and bottom rows, respectively. Orange and blue lines are the average spectra of (subsequent) hit and miss trials, respectively. vertical gray shaded regions indicate p<.05 for the difference between hit and miss spectra after cluster correction. Colored shaded regions indicate the standard error of the mean. Successful encoding elicited greater power in the Hip between 2.0 and 2.9 Hz. Failed retrieval elicited greater power in the PHG between 4.4 Hz and 15 Hz. Although other ROIs did not exhibit hit/miss differences, power spectra nevertheless appeared different across regions and between encoding and retrieval. These differences were explored using linear mixed effects modeling of power as a function of frequency (2-80 Hz; modeled with 8 splines), region, and encoding/retrieval. All interactions between fixed effects were significant (2(4-32)>66,maximum p <2e-10). Several factors may explain these interactions. First, an interaction between frequency and region may have been driven by regional variations in the frequency of maximum power in the lower range (2-20 Hz; Table 1). Second, an interaction between encode/retrieve and region may have been driven by higher power in the Hip during retrieval than during encoding. Third, a three-way interaction may have been driven by secondary power increases between 24 and 32 Hz in the Hip, ACC, and dlPFC during retrieval (Table 1). These results emphasize that HFB peaks constituted physiological events, and not statistically extreme values. D. Similar to A, except time on the x axis is represented in seconds relative to the image onset. E. Similar to D, except for retrieval. F. Heatmaps display the mean difference in z-scored power between hit and miss trials. For all panels, the x-axis displays time relative to image onset, and the y-axis displays frequency. White outlined areas indicate p<.05 for the difference between hit and miss power after cluster correction. In the Hip and dlPFC, there were positive subsequent memory effects in 2 to 10 Hz power just before and during image onset (-450 to 100 ms). In the PHG, there was a negative memory effect in 2 to 10 Hz later after image onset at both encoding (550 to 1550 ms) and retrieval (925 to 1450 ms). These decreases partially overlapped with the visual response observed in the PHG's HFB power (Supplemental Figure 1A). Finally, the pPFC exhibited a late (1225 to 2775 ms) positive subsequent memory effect during encoding across 2 to 10 Hz. At retrieval, the pPFC exhibited a much earlier (-100 to 600 ms) positive memory effect and a later (950 to 1975 ms) negative memory effect.

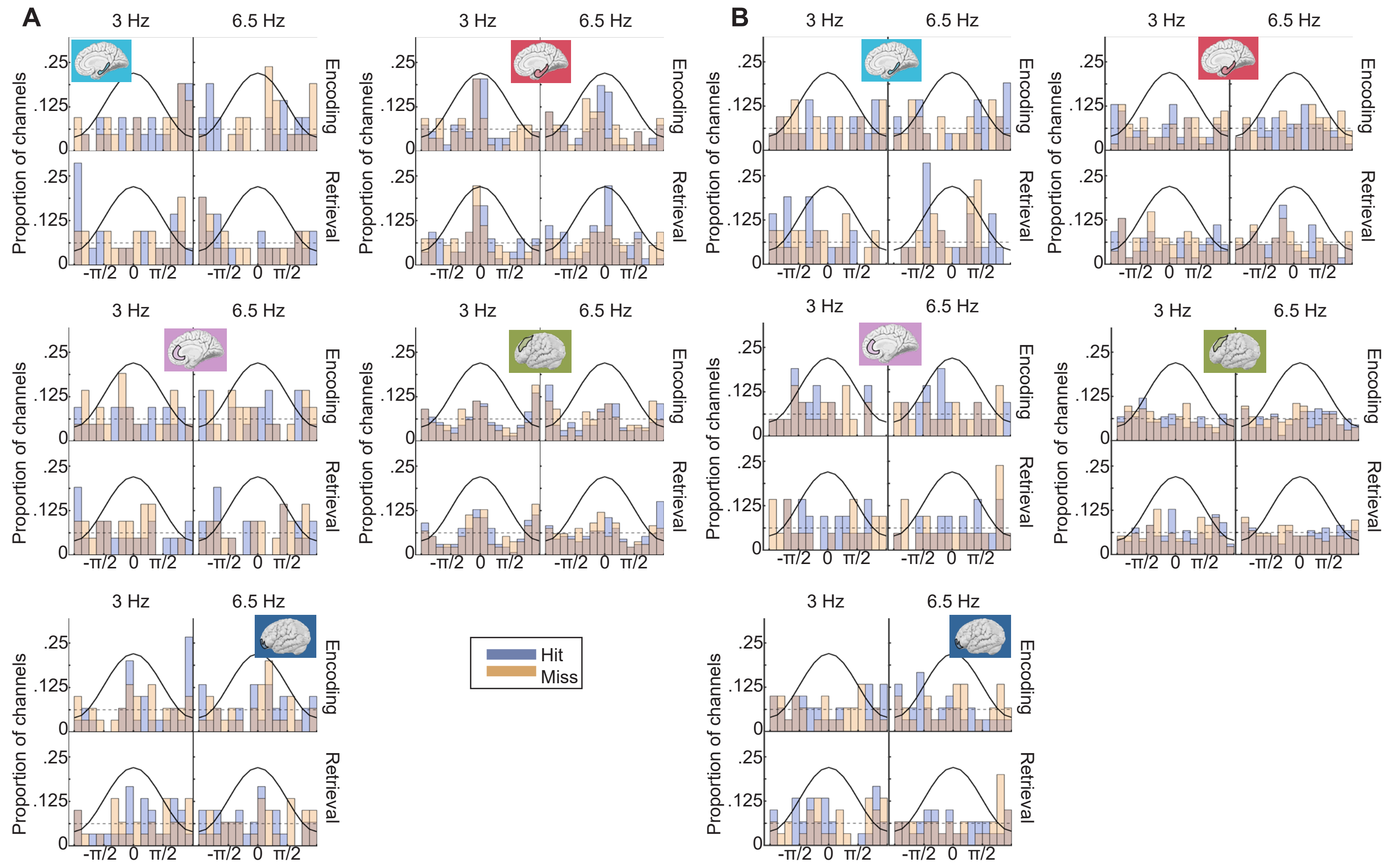


Supplemental Figure 4. Distributions of phase preference of phase clustering within channels. A. Histograms display the proportion of channels on the y axis and the phase of either the 3 Hz (left columns) or 6.5 Hz (right columns) component of the signal. For each channel, the mean circular phase was calculated across trials at the time point of the HFB peak for hit (blue) and miss (orange) trials separately. The superimposed sin wave on each plot indicates which phase values correspond to the peak, trough, or intermediate positions in the oscillation. The dashed dotted lines indicate uniform distribution. The top row of each panel displays encoding data. The bottom row displays retrieval data. Each panel displays data for a different region. Notice that the Hip exhibited a preference for HFB activity to occur at the trough of the 3 Hz oscillation during successful memory trials at both encoding and retrieval, but the PHG exhibited a preference for the peak. B. Similar to A, but mean phase was measured at the time point of maximal ITPC relative to image onset. Notice that phase preferences evident in MTL regions in C appear weaker when measured relative to image onset.

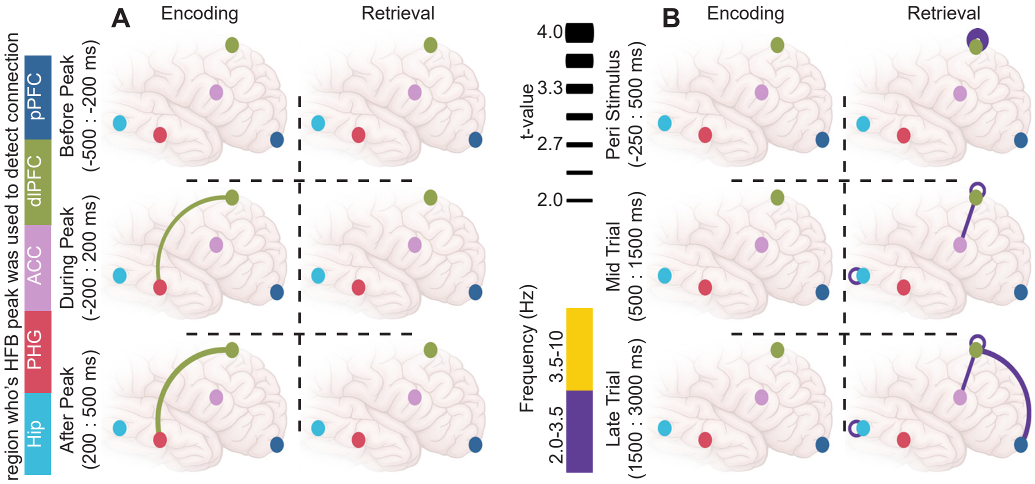


Supplemental Figure 5. Negative mnemonic connections. Plots A and B correspond to main text Figure 4 E and F, respectively. However, while the schematics in Figure 4 E and F display only connections which exhibited increased strength for hit trials, this figure displays only connections which exhibited decreased connectivity strength for hit trials relative to miss trials. Note that far fewer of these negative connections were detected than the positive connections shown in Figure 4.

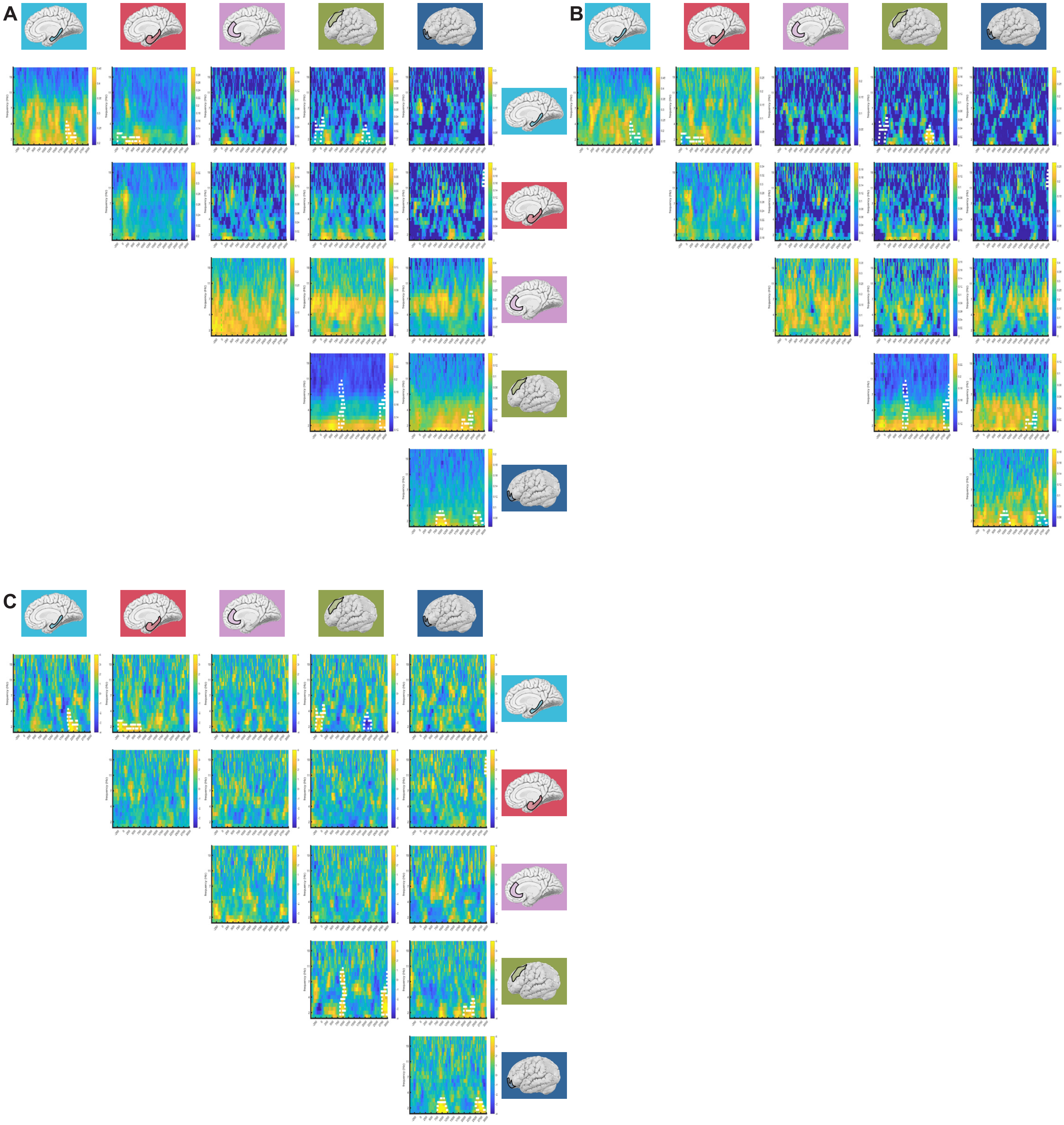


Supplemental Figure 6. Inter regional connectivity changes associated with memory during encoding. Analysis aligned to image onset. In all panels, subplots have time relative to image onset (t=0) on the X-axis and frequency on the Y-axis. The color scale indicates the strength of connectivity between the regions indicated by the row and column of the subplot. A. Connectivity during subsequent hit trials using pairwise phase consistency. B. Connectivity during subsequent miss trials measured using pairwise phase consistency. C. The difference between hit and miss trials. Warm colors indicate stronger connectivity during hit trials. Color scale represents t-values. White outlines indicate cluster-corrected significant differences between hit and miss trials.

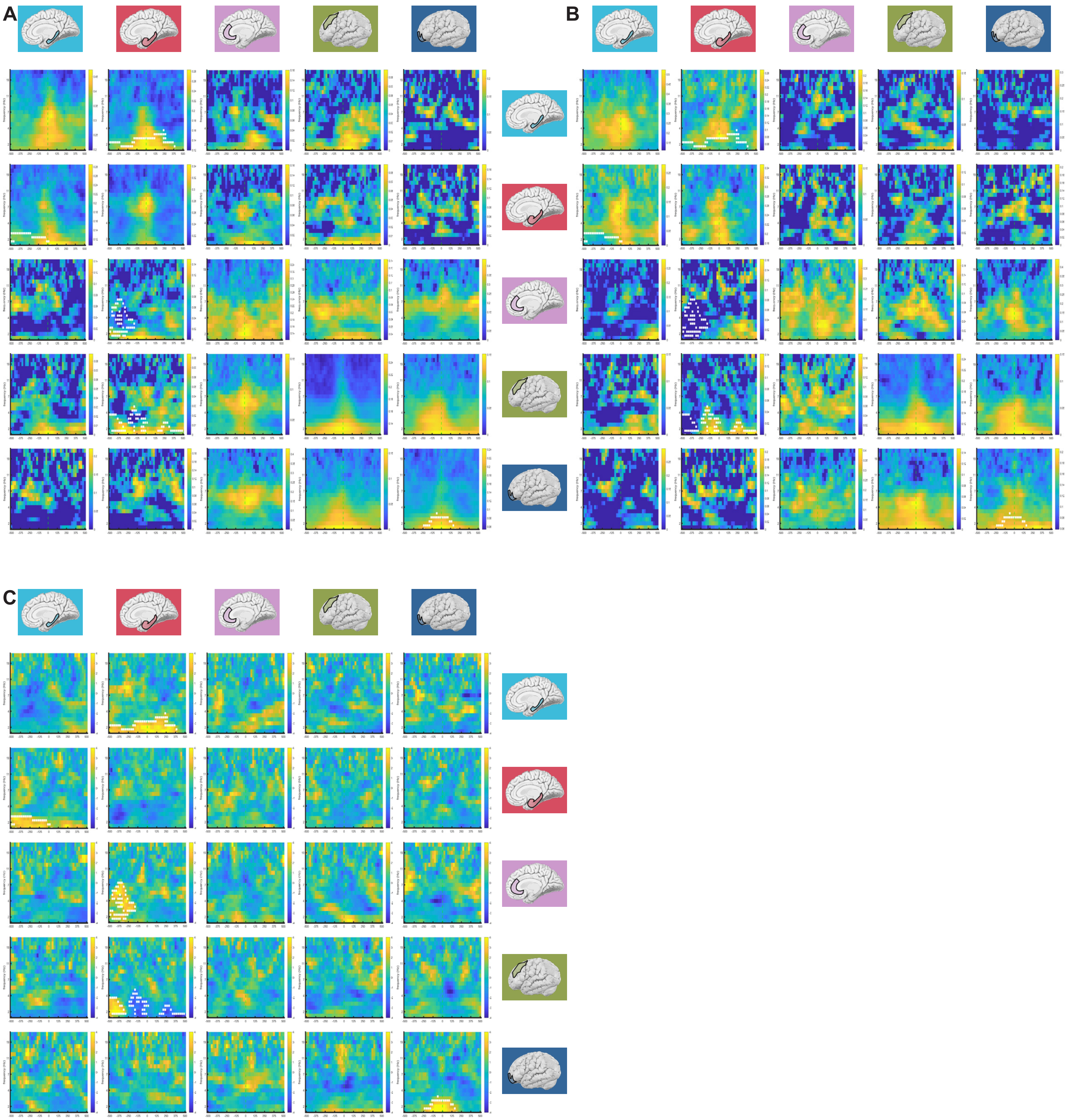


Supplemental Figure 7. Inter regional connectivity changes associated with memory during encoding. Analysis aligned to HFB peak of the region indicated by the row position of each subplot. In all panels, subplots have time relative to HFB peak (t=0) on the X-axis and frequency on the Y-axis. The color scale indicates the strength of connectivity between the regions indicated by the row and column of the subplot. A. Connectivity during subsequent hit trials measured using pairwise phase consistency. B. Connectivity during subsequent miss trials measured using pairwise phase consistency. C. The difference between hit and miss trials. Warm colors indicate stronger connectivity during hit trials. Color scale represents t-values. White outlines indicate cluster-corrected significant differences between hit and miss trials.

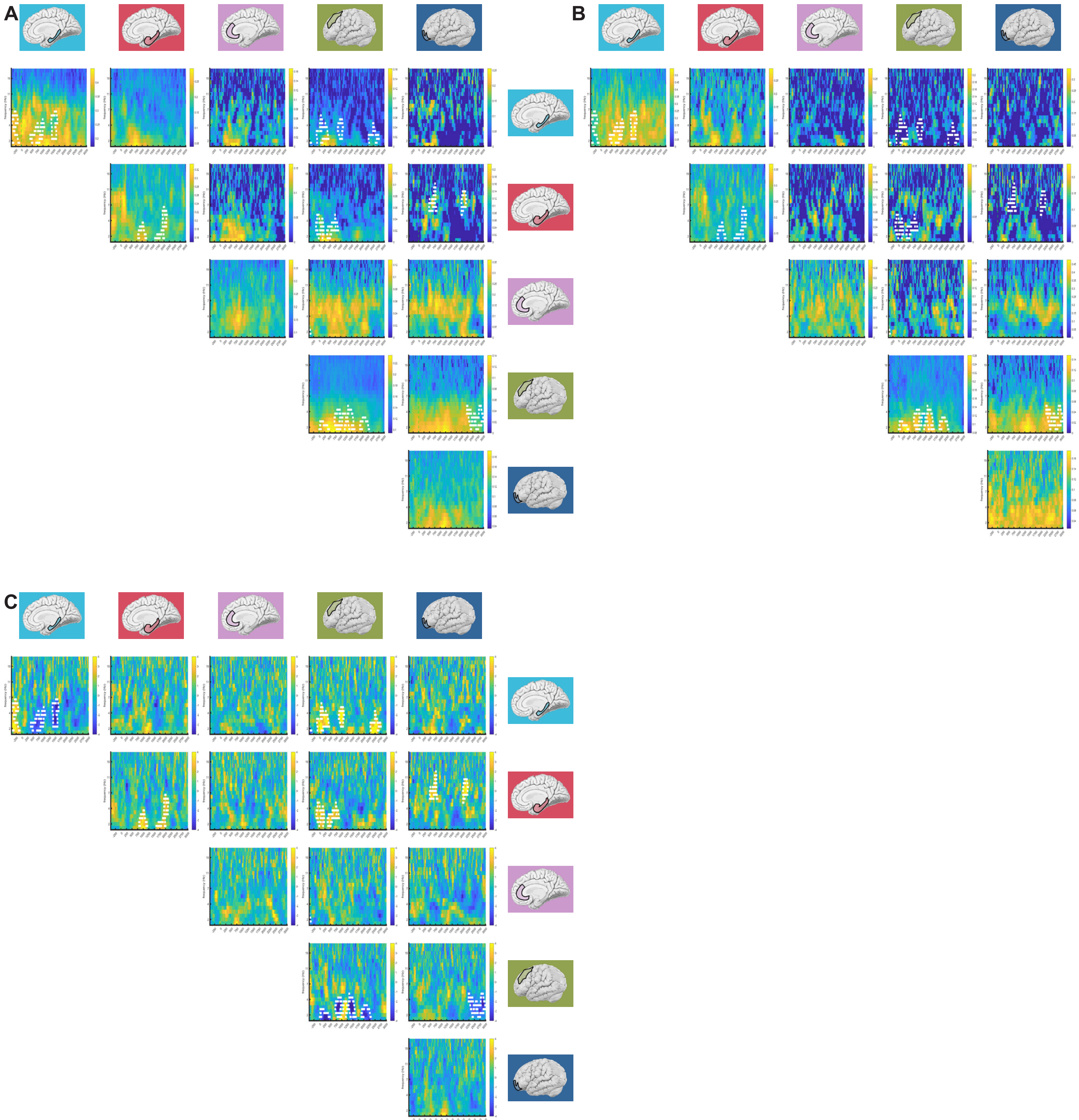


Supplemental Figure 8. Inter regional connectivity changes associated with memory during retrieval. Analysis aligned to image onset. In all panels, subplots have time relative to image onset (t=0) on the X-axis and frequency on the Y-axis. The color scale indicates the strength of connectivity between the regions indicated by the row and column of the subplot. A. Connectivity during hit trials measured using pairwise phase consistency. B. Connectivity during miss trials measured using pairwise phase consistency. C. The difference between hit and miss trials. Warm colors indicate stronger connectivity during hit trials. Color scale represents t-values. White outlines indicate cluster-corrected significant differences between hit and miss trials.

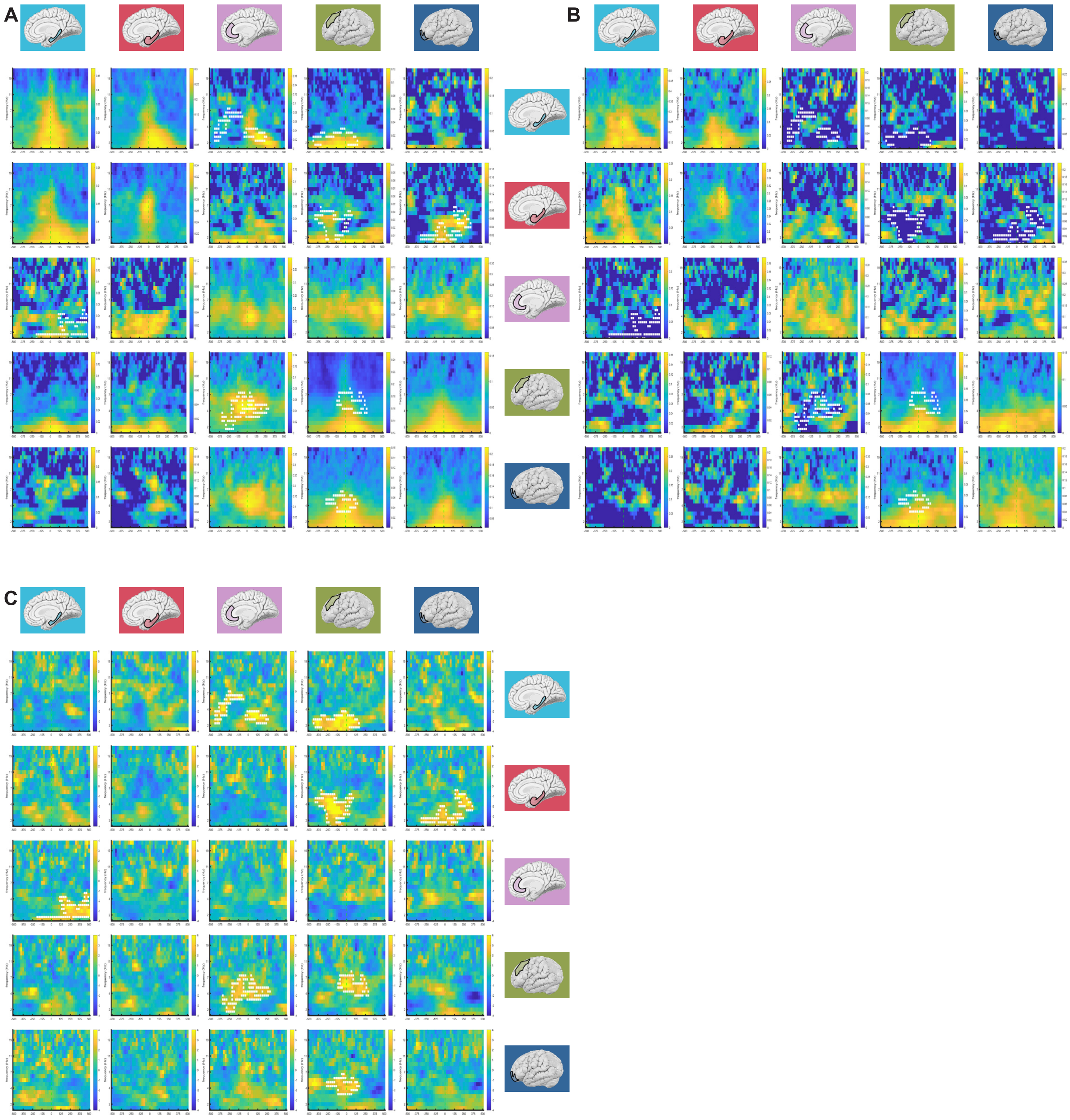


Supplemental Figure 9. Inter regional connectivity changes associated with memory during retrieval. Analysis aligned to HFB peak of the region indicated by the row position of each subplot. In all panels, subplots have time relative to HFB peak (t=0) on the X-axis and frequency on the Y-axis. The color scale indicates the strength of connectivity between the regions indicated by the row and column of the subplot. A. Connectivity during hit trials measured using pairwise phase consistency. B. Connectivity during miss trials measured using pairwise phase consistency. C. The difference between hit and miss trials. Warm colors indicate stronger connectivity during hit trials. Color scale represents t-values. White outlines indicate cluster-corrected significant differences between hit and miss trials.

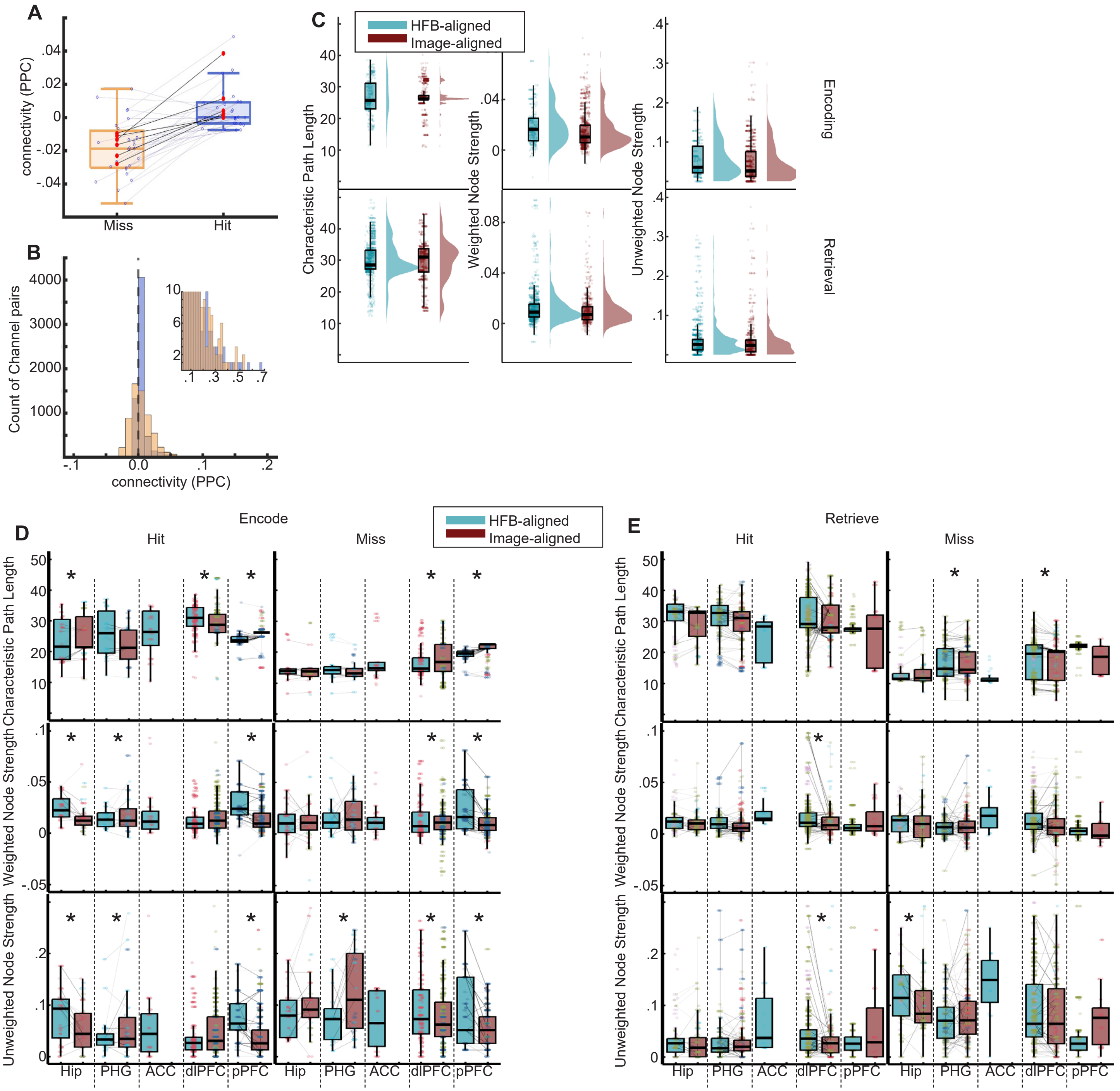


Supplemental Figure 10. Additional graph analysis results. A. Box plot display connectivity between the ACC and Hip for all channel pairs that spanned the two regions (open blue circles) and participants (red circles) averaged across the same time and frequency windows as in Figure 5A.i and A.ii. Note that although miss trials were characterized by more densely connected overall graphs (A.ii), connectivity between the ACC and Hip was weaker during miss trials for all participants. B. The histogram displays the strength of all connections for the representative participant shown in Figure 5A averaged across the long temporal epoch (corresponding to Figure A.iii and A.iv). Note that with lower variance in the hit distribution, a higher central tendency in the hit distribution, and similar representation at very strong connectivity strengths between hit and miss trials (see inset), a z-scored analysis would replicate prior findings of stronger connectivity for hit trials 31. Overall, for this example participant, 52% of all possible connections exhibited greater connectivity values for successful encoding over failed encoding, and the corresponding value was 54% for retrieval trails. It is interesting that overall connectivity was stronger during hit trials when examined over the longer temporal window. C. Same as in Figure 5B but with data sorted by whether significant connections were detected in HFB-aligned or image-aligned analyses. There was a modest effect such that graphs exhibited more connectivity for HFB-locked analyses than image-locked: shorter characteristic path length, increased weighted strength, and increased unweighted strength; 2(1)>12,maximum $p < .0005$. D. Boxplots display results of graph analysis for each region and timeset combination separately during encoding. Top, middle, and bottom panels display characteristic path length, weighted node strength, and unweighted node strength respectively. The colors of dots represent the region of each connection partner. Asterisks indicate significant ($p<.0001$) holm-corrected comparisons between HFB-aligned and image-aligned values within region. Notice that Hip and pPFC graphs are more connected when aligned to HFB peaks during successful encoding. This is true for the pPFC during failed encoding as well. E. Similar to D except for retrieval.

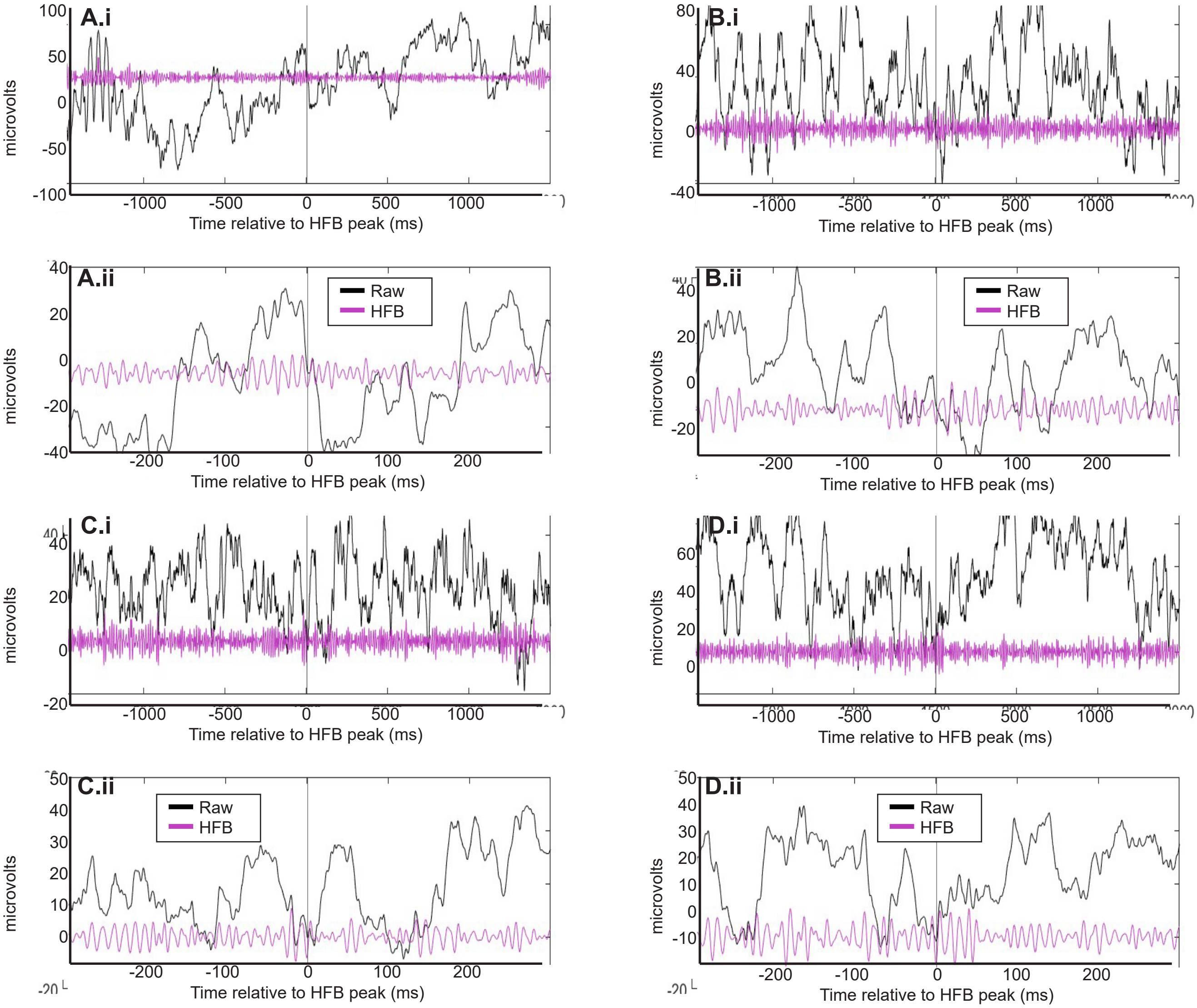


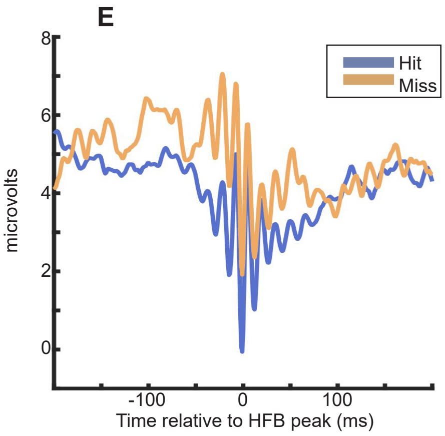


Supplemental Figure 11. Example HFB peaks do not look like sharp wave ripples, but their mean does. A-D Line plots display individual HFB peak events. The raw timeseries is displayed in black and the timeseries bandpass filtered to between 70 and 150 Hz is displayed in purple. Panel ii displays a zoomed in time scale relative to panel i. Time is displayed on the x axis relative to the time point of HFB peak detection. The y axis displays the electrical potential measured in microvolts. All randomly chosen events are from hippocampal channels. Each event is recorded in a different participant. E. The line plot displays the mean of the raw hippocampal encoding timeseries aligned to the nearest trough of the HFB signal relative to the peak of the HFB power for hit and miss trials separately. Notice that although no individual events bear visual resemblance to classical sharp wave ripples, these mean events do.
